# Genome-wide transposon mutagenesis of paramyxoviruses reveals constraints on genomic plasticity

**DOI:** 10.1101/2020.03.30.016493

**Authors:** S. Ikegame, S. M. Beaty, C. Stevens, S. T. Won, A. Park, D. Sachs, P. Hong, P. A. Thibault, B. Lee

**Affiliations:** Department of Microbiology at the Icahn School of Medicine at Mount Sinai, New York, NY 10029, USA; Department of Genetics and Genomic Sciences at the Icahn School of Medicine at Mount Sinai, New York, NY 10029, USA

**Author notes:** These authors contributed equally to this work (S. Ikegame, S. M. Beaty).

## Abstract

The antigenic and genomic stability of paramyxoviruses remains a mystery. Here, we evaluate the genetic plasticity of Sendai virus (SeV) and mumps virus (MuV), sialic acid-using paramyxoviruses that infect mammals from two *Paramyxoviridae* subfamilies (*Orthoparamyxovirinae* and *Rubulavirinae*). We performed saturating whole-genome transposon insertional mutagenesis, and identified important commonalities: disordered regions in the N and P genes near the 3’ genomic end were more tolerant to insertional disruptions; but the envelope glycoproteins were not, highlighting structural constraints that contribute to the restricted antigenic drift in paramyxoviruses. Nonetheless, when we applied our strategy to a fusion-defective Newcastle disease virus (*Avulavirinae* subfamily), we could select for F-revertants and other insertants in the 5’ end of the genome. Our genome-wide interrogation of representative paramyxovirus genomes from all three *Paramyxoviridae* subfamilies provides a family-wide context in which to explore specific variations within and among paramyxovirus genera and species.

## Introduction

The *Paramyxoviridae* family encompasses a diverse and ever-expanding range of mammalian pathogens, including such familiar human viruses as measles (MeV), mumps (MuV), parainfluenza, and henipaviruses (*1*). Paramyxoviruses (PMVs) are negative-sense, single-stranded RNA viruses with genes coding for at least six major proteins: nucleocapsid (N), phosphoprotein (P), matrix (M), fusion glycoprotein (F), receptor binding protein (RBP, formerly designated variously as HN, H, or G), and the large protein (L) that possesses RNA-dependent RNA polymerase (RdRp) activity (*2*). In addition, different virus species encode a host of accessory proteins from the P gene. Others have additional less well-characterized genes (*e.g*. for small-hydrophobic (SH) proteins in some orthorubulaviruses). Despite persisting in human populations for centuries, individual PMVs show a remarkable lack of antigenic variability within the common envelope glycoproteins (F and HN/H/G), and often cross-react to antibodies raised against closely-related viruses (*3–5*). For example, the MeV and MuV strains used in the MMR vaccines have not changed in the last 40 years, and yet are still protective against current field isolates (*6*). Indeed, a MuV-like virus isolated from bats (*7*) is cross-neutralized by mumps-vaccinated human sera (*8*), and the latest ICTV classification considers this bat mumps virus as a strain of MuV rather than a new *Orthorubulavirus* species (*2*). This is in contrast to the well-known propensity for antigenic drift of influenza virus, another negative-sense RNA virus, in response to various pressures including host populations’ adaptive immune responses (*9*).

Our lab previously examined the overall genetic plasticity of a vaccine-strain MeV through whole-genome transposon insertional mutagenesis (*10*), and found that unlike influenza virus (*11*), MeV did not tolerate insertional changes in its surface glycoproteins, F and H. MeV also demonstrated a greater overall intolerance for mutagenesis throughout its genome, concomitant with the known increased genetic stability of PMVs (*3, 12*). However, MeV-H utilizes protein receptors to mediate virus entry (*13*), while a wide range of other PMVs use sialic acids (SAs) to facilitate entry (*14*), like influenza does. Thus, we sought to learn whether divergent SA-using PMVs would demonstrate a tolerance in their attachment glycoproteins that correlated with the virus family, or with receptor usage, and whether we would observe other genetic constraints that were similar to those found in MeV.

We first generated genome-wide transposon insertional mutagenesis libraries of SeV (genus *Respirovirus*) and MuV (genus *Orthorubulavirus*) as representative members from the two subfamilies of *Paramyxoviridae, Orthoparamyxovirinae* and *Rubulavirinae*, that infect mammals. We then evaluated enrichment of transposon abundance across the genome during serial passaging to identify genetic plasticity—the ability to tolerate transposon insertions without a loss of replicative fitness—at any given loci. We found that SeV and MuV show similar trends in genetic plasticity, permitting insertions in the 3’-most N and P genes, and especially in the non-coding untranslated regions (UTRs). We then rescued representative insertion mutants (insertants) among the most-enriched regions to determine the veracity of our transposon mutagenesis library screens. In general, insertants in the N and P genes of both viruses were viable while those elsewhere in the genome were not. Interestingly, in “multiplex” competition assays of viable insertants, we found that SeV insertants demonstrated a differential fitness hierarchy from what was observed in the library setting. In contrast, the fitness of MuV insertants was overall consistent with their enrichment from the original transposon library. Finally, to determine if our experimental strategy had the power to select for mutants and/or insertants that are vanishingly unlikely to occur naturally, that is, in an otherwise inaccessible fitness landscape, we generated a transposon mutagenesis library in a Newcastle disease virus (NDV) background made fusion-defective by a point mutant in its fusion peptide (NDV^Fmut^). Serial passaging of NDV^Fmut^ enriched for F-insertants in the original NDV^Fmut^ background that restored fusion. Interestingly, we also observed an enrichment for an L-insertant that also carried the reversion mutation restoring the functionality of the fusion peptide. Together with our previous studies on MeV, we show that the genetic plasticity of PMVs is broadly consistent across different genera regardless of whether the PMVs use sialic acid-based or protein-based receptors. We also show that our experimental strategy can be used to access and interrogate arbitrarily distant fitness landscapes. Finally, we identify common insertion-tolerant regions within the PMV genomes that can be exploited for engineering recombinant PMVs.

## Results

### Sendai virus broadly tolerates insertions in the 3’ end of the genome

In order to identify which regions of the SeV genome can best accommodate insertional mutations, we utilized a *Mu*-transposon insertional mutagenesis strategy (Fig. 1A) to introduce 15 nucleotide (nt) insertions throughout the SeV genome, which is equivalent to 5 amino acid (aa) insertions if the transposon lands in an open reading frame (ORF). The insertional mutagenesis library approach is a more disruptive approach to interrogating the PMV genome, in comparison to single-nucleotide mutagenesis approaches that are more suited to interrogating single genes. However, insertants induce a severe selective pressure on the virus, which is helpful for whole-genome interrogation. Briefly, we first generated a SeV genomic plasmid with an extra 3-nt stop codon added at the end of the EGFP reporter gene (SeV 6n+3). Since PMV genomes follow the “rule-of-six”, where the entire genome length must be an exact multiple of six (6n) in order to replicate well (*15*), SeV 6n+3 should be rescued inefficiently and replicate even less so on its own (Fig. 1B and data not shown). The same is true for MuV, whose genome we also interrogated similarly in this study (Fig. 1C). Since our intention was to understand the tolerance of each of the virus’ native genetic regions for insertions, we then excluded the reporter gene from all downstream analyses. We subjected this SeV 6n+3 plasmid (‘parental 6n+3’ in Fig. 1A) to *Mu*-transposon mutagenesis using optimized conditions to achieve saturating mutagenesis, ultimately leaving random 15-nt insertions across the genome. Transposon-mutated genomic plasmid libraries are therefore 6n+18 (Fig. 1A), restoring the “rule-of-six”, and should be more competent for rescue and replication (Fig. 1B, C). In addition, any genomes that failed to receive the transposon remain 6n+3 in length; these genomes cannot be rescued well, and ultimately will not be represented in our sequencing analyses because they lack the transposon sequence. Importantly, the 15-nt transposon ‘scar’ itself is designed to be translatable in all three reading frames. The SeV 6n+18 insertional library was rescued in BSRT7 cells with rescue events (~3 x 10^5^) equal to approximately 19-fold coverage of the genome (Supplementary Table S1). Rescued virus from the supernatant was passaged twice in biological triplicates at a low multiplicity of infection (moi) on Vero cells, which should screen for the replicative fitness of any given insertion. We chose Vero cells – lacking interferon signaling – as a neutral background to remove confounding antiviral selective pressures in our experiments. In particular, the P gene encodes the phosphoprotein, an essential co-factor for the viral transcriptase and replicase, but also encodes accessory proteins to mitigate the host interferon response. In the absence of interferon signaling, insertants that disrupt accessory proteins are not selected against and we can better explore the structural and functional plasticity of the phosphoprotein itself. Thereafter, the original library plasmid pool (input), supernatant from the rescue (P0), and passages (P1, P2) were deep-sequenced and evaluated for the presence of the 15-nt insertion.

**Figure 1.**
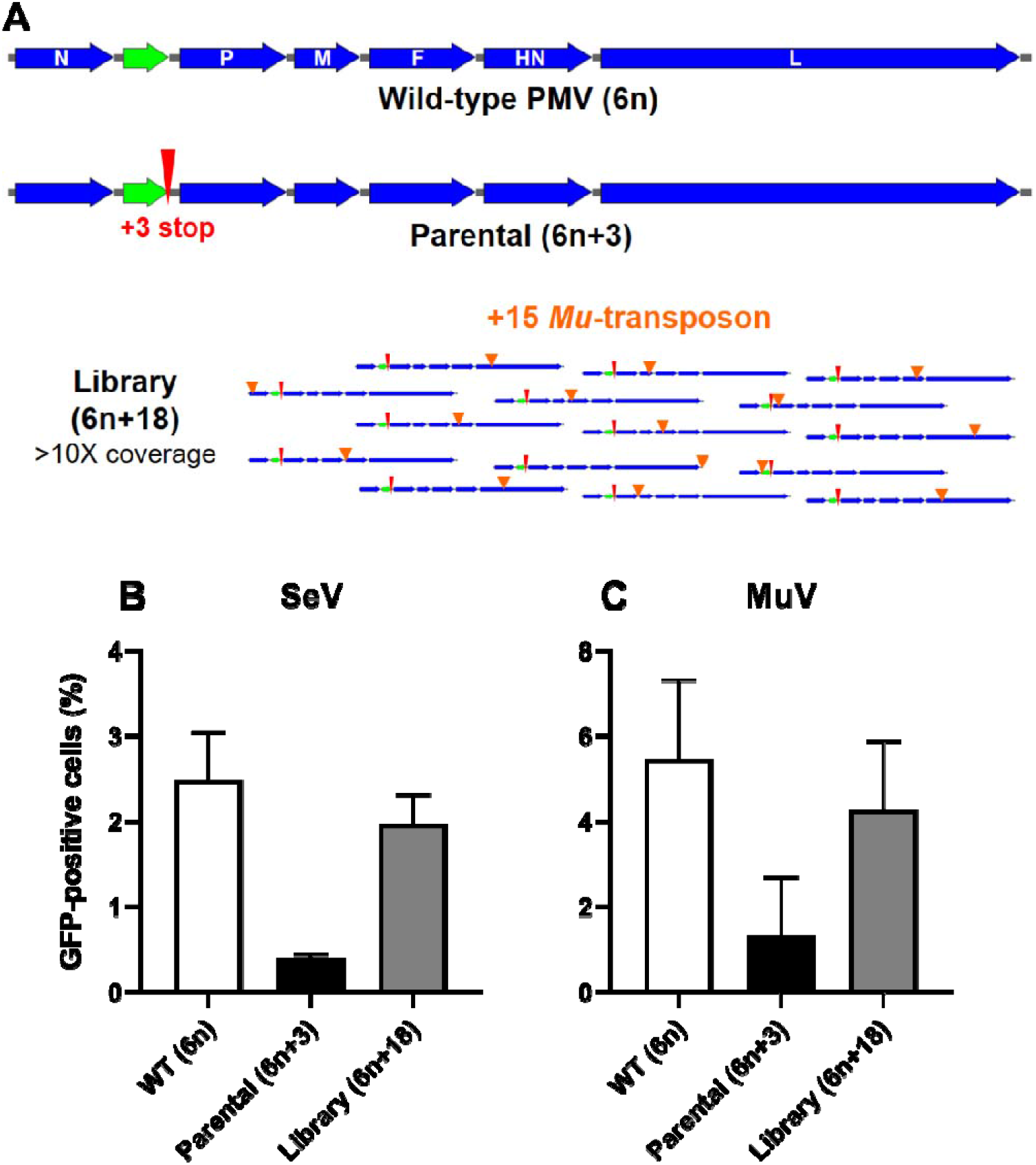
Insertional mutagenesis strategy. (A) Generation and selection of our transposon mutagenesis library. Schematic of a generic paramyxovirus genome that conforms to rule-of-six. Wild-type PMV (6n), is shown in deep blue with the additional EGFP reporter between the N and P genes shown in green. The parental (6n+3) genome used for generating the transposon mutagenesis library is shown below the wild-type PMV (6n) genome. Red elongated arrowhead indicates the additional stop codon (+3 stop). This 6n+3 genome should be rescued much less efficiently than the wild-type 6n genome. Library (6n+18) shows examples of the random +15 (nt) *Mu*-transposon scar (light blue arrowheads) left in the 6n+3 parental genome after library generation. Each library was generated at a scale to result in >10X coverage (number of individually-rescued insertants), in order to ensure that every nt position has >90% probability of having at least one transposon insertion, assuming *Mu*-transposon insertions are random. **(B-C) Relative rescue efficiencies of SeV (B) and MuV (C) constructs.** WT (6n), Parental (6n+3), and the transposon mutagenesis Library (6n+18) genomes from SeV and MuV were generated and rescued as described in Materials and Methods. Relative rescue efficiencies were estimated by FACS analysis and indicated on the y-axis as GFP-positive cells (%) at 48 hpi as detailed in Materials and Methods.

In the input plasmid pool, insertions were found at 65.9% of nucleotide sites, and ultimately targeted ~93% of amino acids throughout the genome (Supplementary Table S1: Library metrics). Importantly, insertions were distributed evenly across the genome in the plasmid pool (Supplementary Fig. S1), demonstrating no bias in the input SeV 6n+18 library.

We mapped insertion coverage from P0, P1, and P2 onto the SeV genome (Fig. 2A-C) and observed a clear purifying selection upon passaging, as expected. At P2, we observe a clear enrichment for insertions in the N and P genes, located at the 3’ end of the genome, and a depletion of insertions elsewhere in the genome. The magnitude and location of insertional enrichment is relatively reproducible between each of the three replicates seen in Fig. 2D. The consistent preference for insertions in the 3’ end of the genome, and particularly the non-coding region between N and P, is also clear. To determine if there were other broad patterns to insertant enrichment, we then analyzed the change in insertional frequency from P0 to P2 in the 5’ UTR, ORF, and 3’ UTR of each viral gene (Fig. 2E). We observed significant enrichment of the 5’UTR-P over its cognate ORF; similar trends can be seen with the other ORFs and their cognate UTRs when they are well-represented at P2. HN and L genes have much smaller UTRs and drastically fewer insertants by P2, making interpretation of their relative enrichments difficult. We also specifically observed that the 5’ UTR of the N gene and 3’ UTR of the L gene both show reduced insertions relative to their neighboring regions, indicative of the stringency of the additional roles these regions play as the 3’ leader and 5’ trailer sequences of the virus, required for viral genomic amplification.

**Figure 2.**
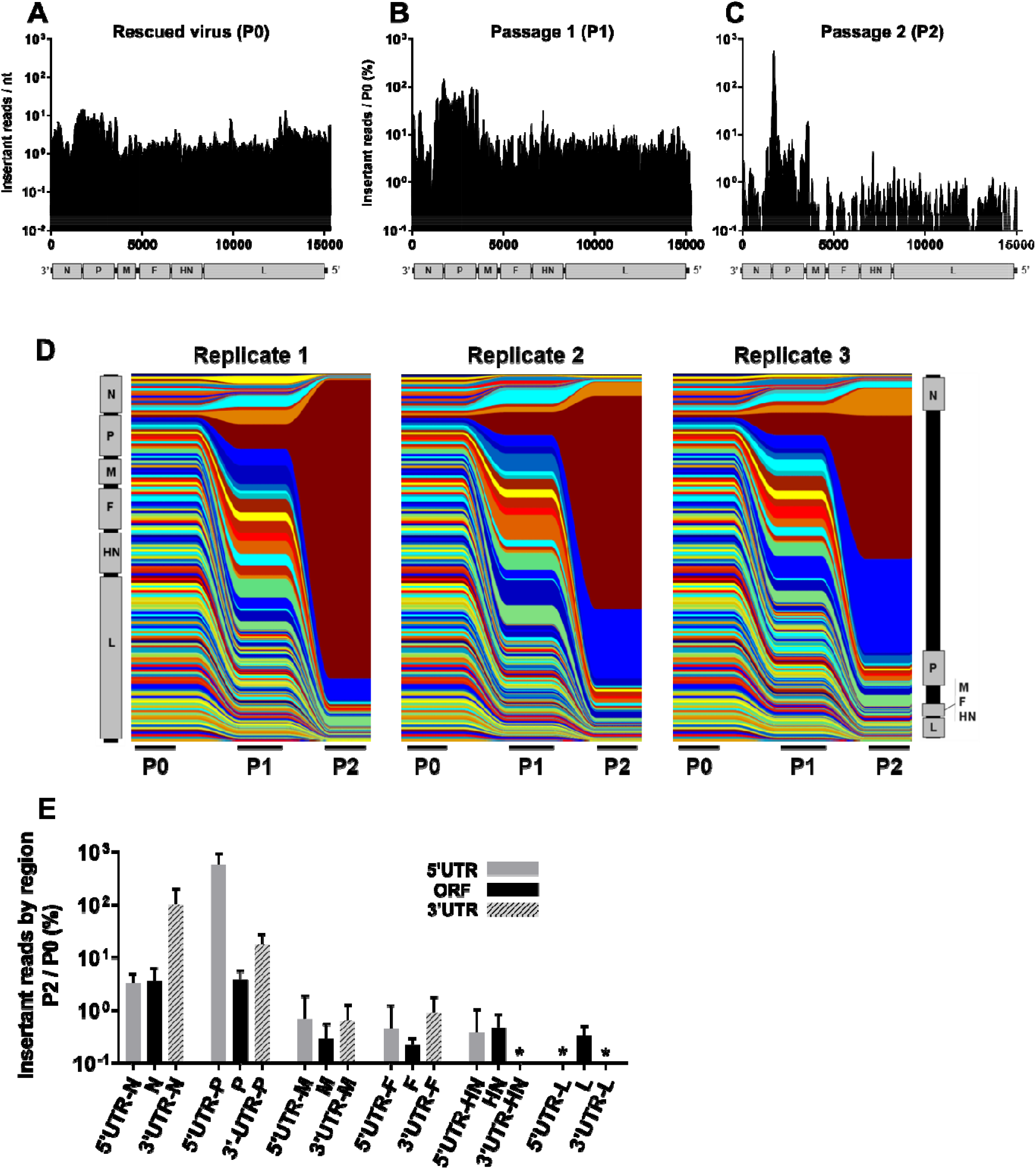
SeV tolerates insertions in non-coding regions and ORFs of highly-expressed genes. Distribution of insertions in a 100nt sliding window identified at P0 (**A**), P1 (**B**), and P2 (**C**). To-scale schematic of the SeV genome is included at the bottom of each graph, which corresponds to the numerical labeling of the genomic nucleotide positions indicated on the x-axis. The y-axis for P0 (A) represents the average number of reads with a transposon scar (insert) (Insertant reads / nt) within a 100-nt sliding window surrounding the genomic nucleotide position indicated on the x-axis. P1 and P2 (B and C, respectively) y-axes are insertant reads / nt as defined in (A) averaged over three biological replicates and expressed as percent of P0 insertant reads / nt, ‘Insertant reads / P0 (%)’. This normalizes the P1 and P2 results for the actual input received from P0. The enrichment or depletion of insertants is therefore a better reflection of the underlying biology and less confounded by the any potential skewing of the input population. (**D**) Stream graph showing the enrichment and/or depletion of insertants over serial passaging (P0, P1, P2) in each of the three biological replicates. Each color represents a sequential 100nt section of genome, with relative abundance at each passage represented by color height. The vertical representation of the SeV genome on the far left and right reflects the distribution of insertants across the genome at P0 and P2, respectively. (**E**) This is a bar graph representation of the P2 data in (C), but separated into the protein coding regions (ORF, black bars) and their respective non-coding regions (5’ and 3’ UTRs, grey and striped bars) as indicated on the x-axis. Data are shown as the normalized averaged ‘Insertant reads in P2 / /P0 (%)’ (y-axis) in each of the ORFs and UTRs. Error bars indicate standard deviation. * indicates values below 0.1%.

### Selected SeV insertants have a different fitness hierarchy in focused competition assays

To evaluate the validity of the results from our pooled rescue and passaging experiments, we re-generated a subset of the most-highly represented SeV 6n+18 single insertants (see Supplementary Table S2). At times, our choices were dictated by cloning successes as some insertants were inexplicably refractory to cloning. To begin, we chose two insertants from each of the most-enriched regions (N-ORF, 3’-UTR-N, 5’-UTR-P, and P-ORF); generally, we chose highly-represented insertants from the original library, but we also chose insertants that are more evenly distributed across those regions of interest. Since some insertants did not rescue (Fig. 3A, discussed below) we included an additional N-ORF-1405 insertant to our panel to maintain representation of highly-enriched regions. Finally, we added the most-enriched insertant for each of the remaining M, F, HN, and L genes, for a total of thirteen insertants (Fig. 3A). We rescued each insertant and amplified it separately, monitoring viral replication kinetics (Fig. 3B, C) and peak titer (Fig. 3A). Two of the insertants (HN-ORF and L-ORF) could not produce infectious virus upon rescue with our highly-efficient reverse genetics system (Fig. 3A, Supplementary Table S2), and a further three insertants (N-ORF-1684, M-ORF, and F-ORF) produced peak titers that were too low for downstream applications (Fig. 3A), indicating that these insertants likely relied on other genomes in the original pool to complement their defects in replication. The remaining eight insertants grew well relative to wild-type SeV, and produced sufficient virus for use in our downstream assay.

**Figure 3.**
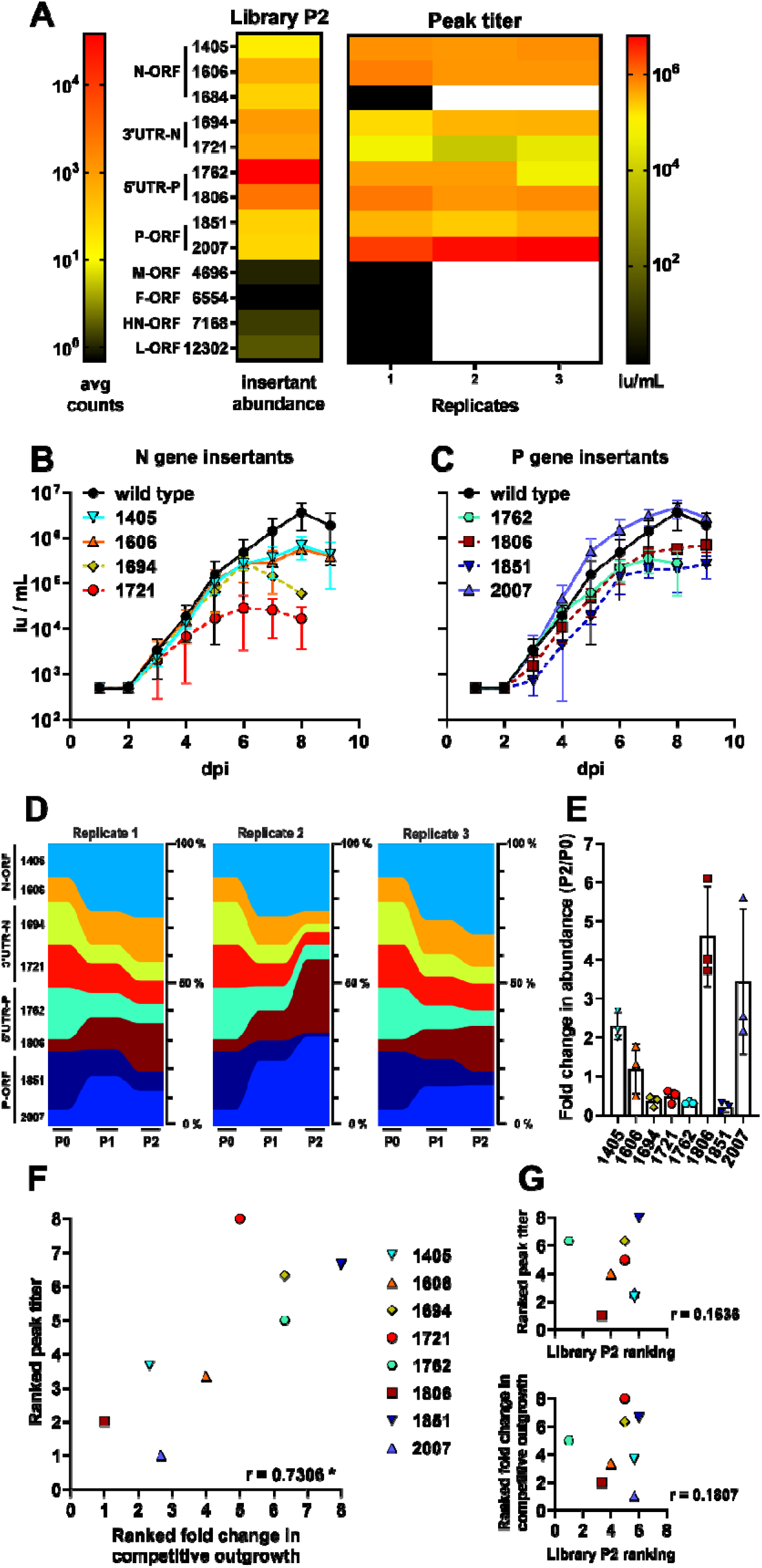
SeV insertant peak titer production predicts fitness in a competitive outgrowth assay. Heat-map comparison of the abundance of select insertants in P2 from our NGS data (left, library P2), and peak titers of highly-represented insertants that were selected for individual confirmation as a recombinant parental virus (6n+3) bearing that particular insertion (+15) (right, Peak titer). Peak titers are indicated from three independent growth curves (Replicates 1, 2, 3). Black blocks in the heat map indicate insertants that failed to produce detectable virus in rescue and so could not be used for any further replicates (indicated by following white blocks). The color-intensity scale for the heat maps comparing the relative abundance of insertants in P2 (avg counts), and the peak titers of selected insertants described above, are indicated on the left and right sides, respectively. (**B** and **C**) Multicycle growth curves of select insertants in the N and P gene regions on Vero cells inoculated with an moi of 0.01. Data are from three independent experiments; mean +/- S.D. are shown as infectious units/ml (iu/ml; y-axis) at the indicated dpi (x-axis). (**D**) Multiplex competitive outgrowth assays on Vero cells, using the insertants characterized from (B) and (C) at a total moi of 0.01. Inoculum and passaging is described in Materials and Methods. Data from three independent replicates are shown as stream graphs: height of color represents the relative abundance (% of total viruses) of the indicated insertants at each passage. **(E)** Bar graph of fold enrichment in % abundance of each insertant from input (P0) to P2 of the competitive outgrowth assay in (D). **(F** and **G)** Comparison of fitness by each of the three major assays by average ranking of the insertants from most fit (1) to least fit (8) in each assay replicate, and their correlation by Spearman non-parametric analysis. * P < 0.05. Ranking is described in Materials and Methods. See the text and Materials and Methods for relevant details.

Next, we sought to evaluate the fitness hierarchy of the insertants. Using a multiplex competition assay that more closely reflects the selective pressures in the passaging library screen, we evaluated if the insertants’ relative abundance in this assay would correlate with either their peak titer or representation in the original library. So, the eight insertants that could be rescued and replicated well, four each from the N and P genes (Fig. 3B-C), were pooled at equal titers and used to infect Vero cells at a total moi of 0.01. We then monitored the fitness of the selected insertants in this competitive outgrowth assay by measuring their expansion over two passages, using MinION long-read sequencing (Fig. 3D, E). To better understand the biological meaning of our library output (*i.e*., insert abundance at P2), which assays predicted each other’s outcomes, and where we ranked our selected insertants relative to each other in each assay, we found that peak titer was tightly-correlated with performance in the competitive outgrowth assay (Fig. 3F), but that ranking from the library screen did not predict either downstream measure of relative fitness (Fig. 3G). This suggests that the SeV n+18 screening library output was predictive of insertant viability – *i.e*. whether it is replication-competent – but not necessarily of relative fitness compared to other genomes. Thereafter, to determine if the complex epistatic interactions we observed in in SeV apply to other sialic acid-using paramyxoviruses, we next turned our analysis to the orthorubulavirus MuV.

### Mumps virus is less tolerant to insertional mutagenesis

Using the same rule-of-six-based strategy as we did with SeV (Fig. 1A, C), we generated a MuV 6n+3 parental genome with which to carry out whole-genome transposon insertional mutagenesis and produced a MuV 6n+18 saturating library of insertants. This library was rescued in BSRT7 cells to ensure a minimum of ten-fold coverage of the genome (>1.6 x 10^5^ rescue events), then passaged twice on Vero cells in biological triplicates as was done for the SeV 6n+18 insertional mutagenesis library. Coverage metrics from deep sequencing are found in Supplementary Table S1 and Supplementary Fig. S2. While nucleotide and codon coverage of the MuV 6n+18 library was similar to that of SeV 6n+18, there was a much larger drop-off in coverage in the viable genomes upon rescue (loss of coverage at 38% and 16% of nt positions in MuV and SeV, respectively, at P0), and an unexpectedly low titer of P0 rescued virus (3.2×10^2^ iu/mL *vs*. 2.2×10^5^ iu/ml for SeV). These initial results suggest that the MuV genome has a much lower overall tolerance for insertions.

### Mumps virus tolerates insertions in the 3’ end of its genome

Due to the low titer from rescue (P0), P1 was carried out at an moi of 0.0001. After two passages, the MuV 6n+18 library also showed evidence of purifying selection (Fig. 4A-C). We observed some enrichment for insertions at the 3’ end of the genome (N and V/P genes) as was seen with SeV, and a surprising secondary peak in the F coding region (Fig. 4C). Stream graphs of each passage replicate (Fig. 4D) show that in comparison with SeV, there was more variability in the regions of the MuV 6n+18 library that were enriched between replicates. This may be a function of the reduced overall coverage of the library, which may allow for stochastic rescue and amplification of viruses that pass a certain viability threshold. Finally, in comparing the specific gene regions that permitted insertions (Fig. 4E), we observed a preference for only certain UTRs over gene coding regions such as the 5’ UTRs of V/P and SH.

**Figure 4.**
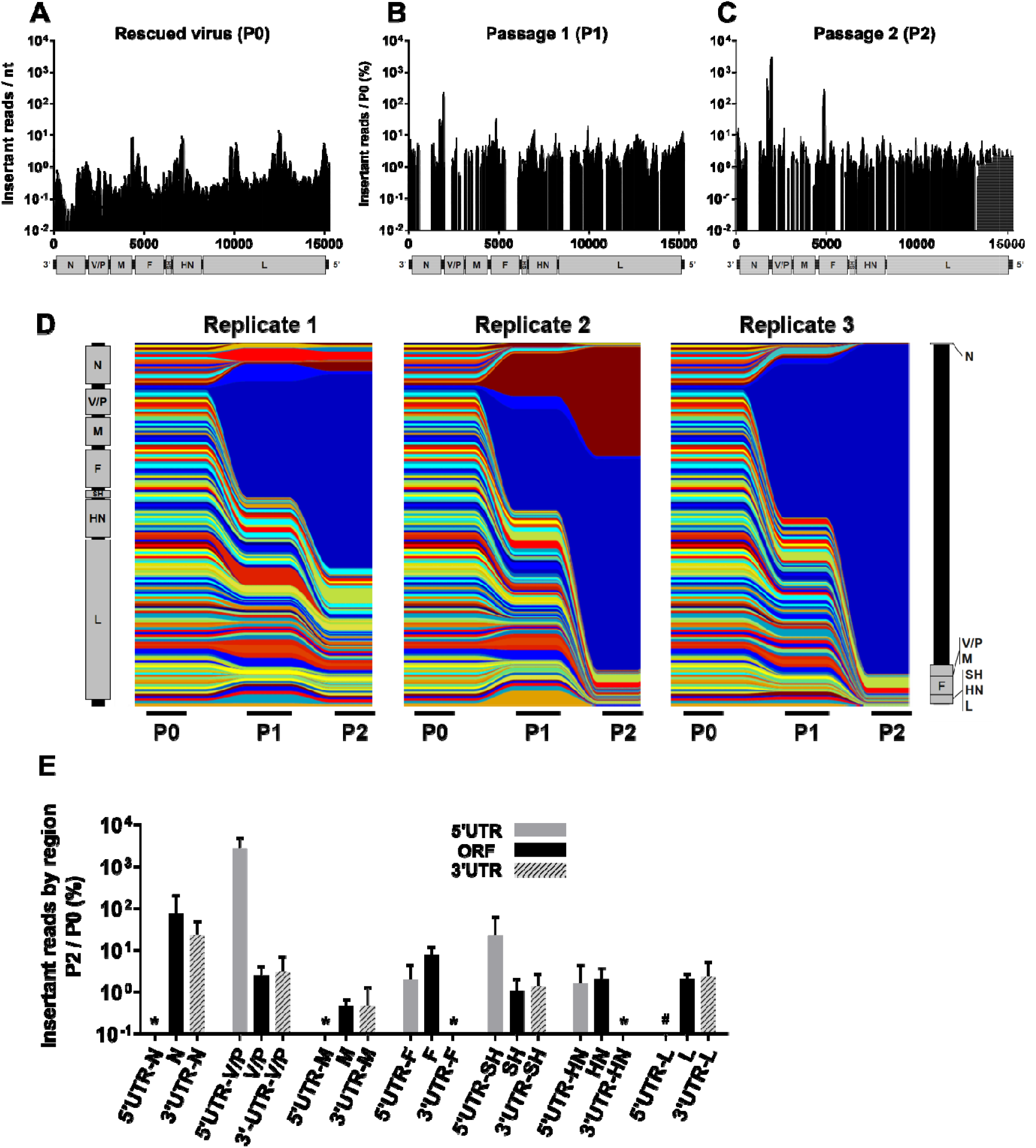
MuV tolerates insertions in non-coding regions and N and F ORFs. Distribution of insertions in a 100nt sliding window identified at P0 (**A**), P1 (**B**), and P2 (**C**). To-scale schematic of the MuV genome is included at the bottom of each graph, which corresponds to the numerical labeling of the genomic nucleotide positions indicated on the x-axis. The y-axis for (A), Insertant reads/nt, and for (B-C), Insertant reads/P0 (%), are defined as in Fig. 2A-C. (**D**) Stream graph showing the enrichment and/or depletion of insertants over serial passaging (P0, P1, P2) in each of the three biological replicates, as described for SeV in Fig. 2D. The vertical representation of the MuV genome on the far left and right reflects the distribution of insertants across the genome at P0 and P2, respectively. (**E**) This is a bar graph representation of the P2 data in (C) but separated into the protein coding regions (ORF, black bars) and their respective non-coding regions (5’ and 3’ UTRs, grey and striped bars) as indicated on the x-axis. Bars represent normalized averaged ‘Insertant Mutant reads in P2/P0 (%)’ +/- S.D. in the indicated genomic regions. * indicates values below 0.1%.

### Mumps virus competitive outgrowth assay reflects threshold viability of enriched insertants in library screen

In order to assess the validity of the MuV 6n+18 library results, we chose the most-highly enriched insertants overall from the library (Supplementary Table S3), representing the N and V/P regions of the genome, as well as the F coding region. Once again, we also included the highest represented insertant from each of the remaining genes (M, HN, and L) in an attempt to more evenly evaluate the library, and rescued each of these insertants. As with SeV, we were unable to rescue the insertants in M, HN, and L (Supplementary Table S3, Fig. 5A), but we were surprised to find that the F insertants also could not be rescued to produce infectious virus. It is likely that these insertants became enriched in the context of the library at P2 by relying on either compensatory mutations or complementation by other genomes. Precedence for the latter is demonstrated by the G264R MeV-F mutant: in the context of an adversely tagged MeV-H where neither wild-type nor G264R MeV-F resulted in syncytia, only viruses with diploid genomes independently bearing the wild-type and G264R MeV-F are able to form syncytia (*16*).

**Figure 5.**
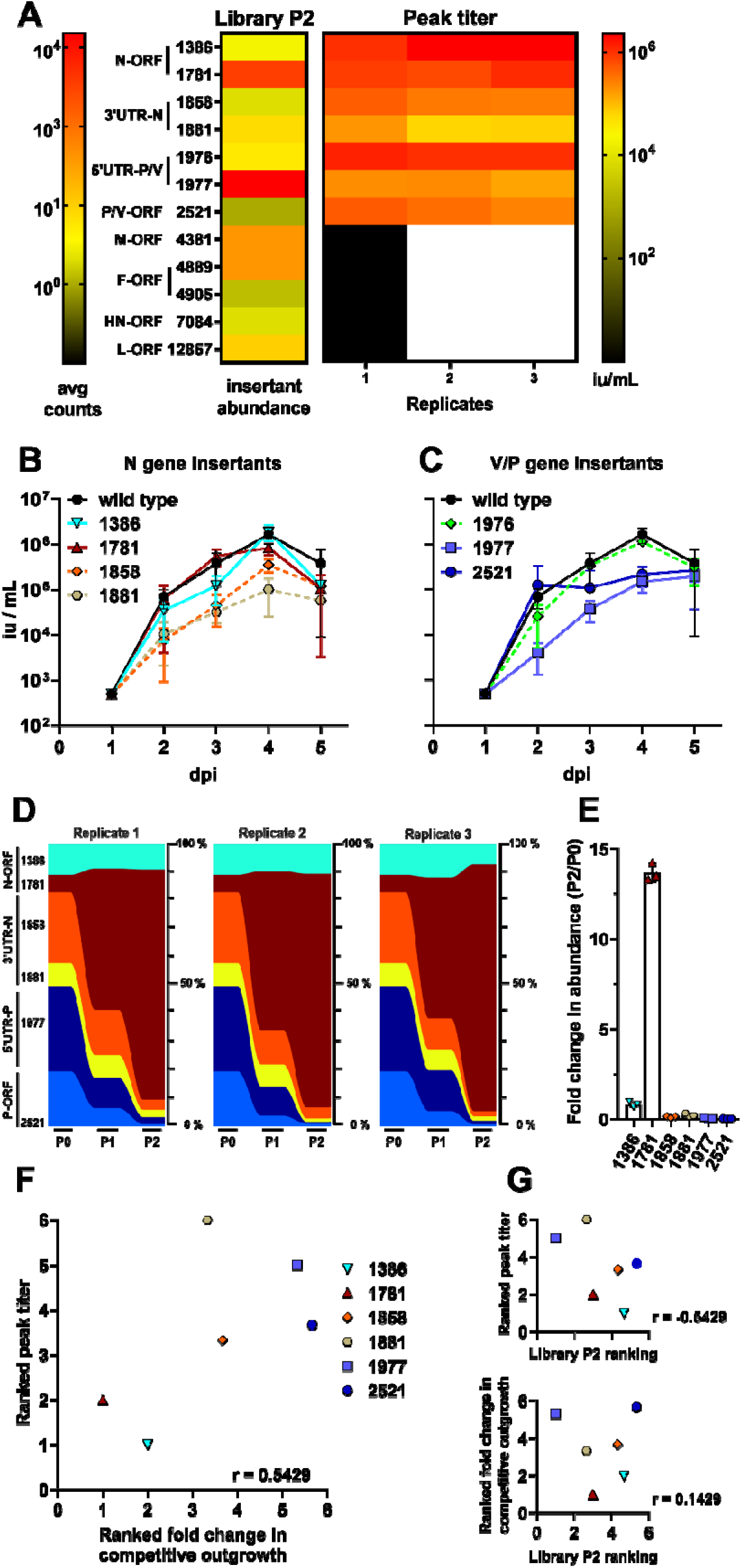
The relative abundance of insertants in the N/P gene regions of MuV from the library P2 correlates with the viability and replicative fitness of those individual insertants. (**A**) Relative abundance of selected insertants in the mutagenesis library after P2 (left, Library P2 column), and their peak titers following rescue and amplification as described in Fig. 3A (right, Peak titer columns), are shown as a heat-map for comparison purposes. The color intensity scale for insertant abundance in Library P2 (avg counts) and the peak titers of the selected insertants are indicated on the left and right sides, respectively. (**B** and **C**) Multicycle growth curves of MuV N-insertants (B) and P-insertants (C) gene regions were generated on Vero cells as described for SeV in Fig. 3B-C. Data are shown as mean +/- S.D. (iu/ml) from three independent experiments at the indicated dpi. (**D**) Multiplex competitive outgrowth assays on Vero cells, using the MuV insertants characterized from (B) and (C), and carried out as described in Materials and Methods. Stream graphs showing the data from three independent replicates are shown: height of each color stream represents the relative abundance (% of total viruses) of the indicated insertants at each passage. **(E)** Bar graph of fold enrichment in % abundance of each insertant from input (P0) to P2 of the competitive outgrowth assay in (D). **(F** and **G)** Comparison of fitness by each of the three major assays by average ranking of the insertants from most fit (1) to least fit (6) in each assay replicate, and their correlation by Spearman non-parametric analysis. Ranking is described in Materials and Methods. See the text and Materials and Methods for relevant details.

Once rescued, we evaluated the remaining successful insertants for growth kinetics and peak titers (Fig. 5A-C), noting that overall these insertants grew well relative to wild-type MuV. We then pooled six of the insertants at equal titers, and infected Vero cells with a total moi of 0.01 in a competitive outgrowth assay. Because the sequencing resolution afforded by the Oxford Nanopore MinION cannot consistently distinguish between insertants P-5’UTR-1976 and −1977, which are shifted by only a single nucleotide, we selected P-5’UTR-1977 for use in the competitive outgrowth assay as it was best-represented in the original library screening (Supplementary Table S3). Over two passages, we evaluated the distribution of the insertants (Fig. 5D, E), and unlike SeV, observed a clear dominance of the N coding region insertants – particularly, N-ORF-1781 – over all the other insertants. The other insertants were only found at 4-40 reads out of the ~1,000 reads in each sample. N-ORF-1386 is a distant second, but still clearly dominant over the other clones. These two insertants also showed the highest peak titer in Fig. 5A-C, along with the excluded P-5’UTR-1976. Evaluating the three assays for correlation by ranking, we determined that peak titer and competitive outgrowth were best correlated (Fig. 5F), while neither correlated well with the original library (Fig. 5G) similarly to what we observed with SeV. Cumulatively, this indicates that while the library screen was valuable for identifying viable insertants, it was not predictive of relative fitness in downstream assays, whereas fitness in one downstream assay predicts relative fitness in another reasonably well.

### Fusion-defective Newcastle disease virus allows access to novel fitness landscapes

In order to (1) test the selective power of our transposon mutagenesis experimental set-up, and (2) determine if the consistent enrichment of insertants in the 3’ end of the genome is a technical artifact of our system, we created a fusion-incompetent NDV by changing a naturally occurring NotI site in the fusion peptide of our NDV genome (see schematic for NDV^Fmut^ in Fig. 6A). Recall that our transposon mutagenesis screen requires that the plasmid encoding the viral genome be free of NotI sites. We hypothesized that the vast majority of insertants in this fusion-defective (NDV^Fmut^) genomic background would not be viable and could not be rescued unless (i) the insertant(s) directly compensated for the fusion-peptide mutation (F^A138T^), and/or (ii) the insertants occurred on a fusion-revertant genome.

**Figure 6.**
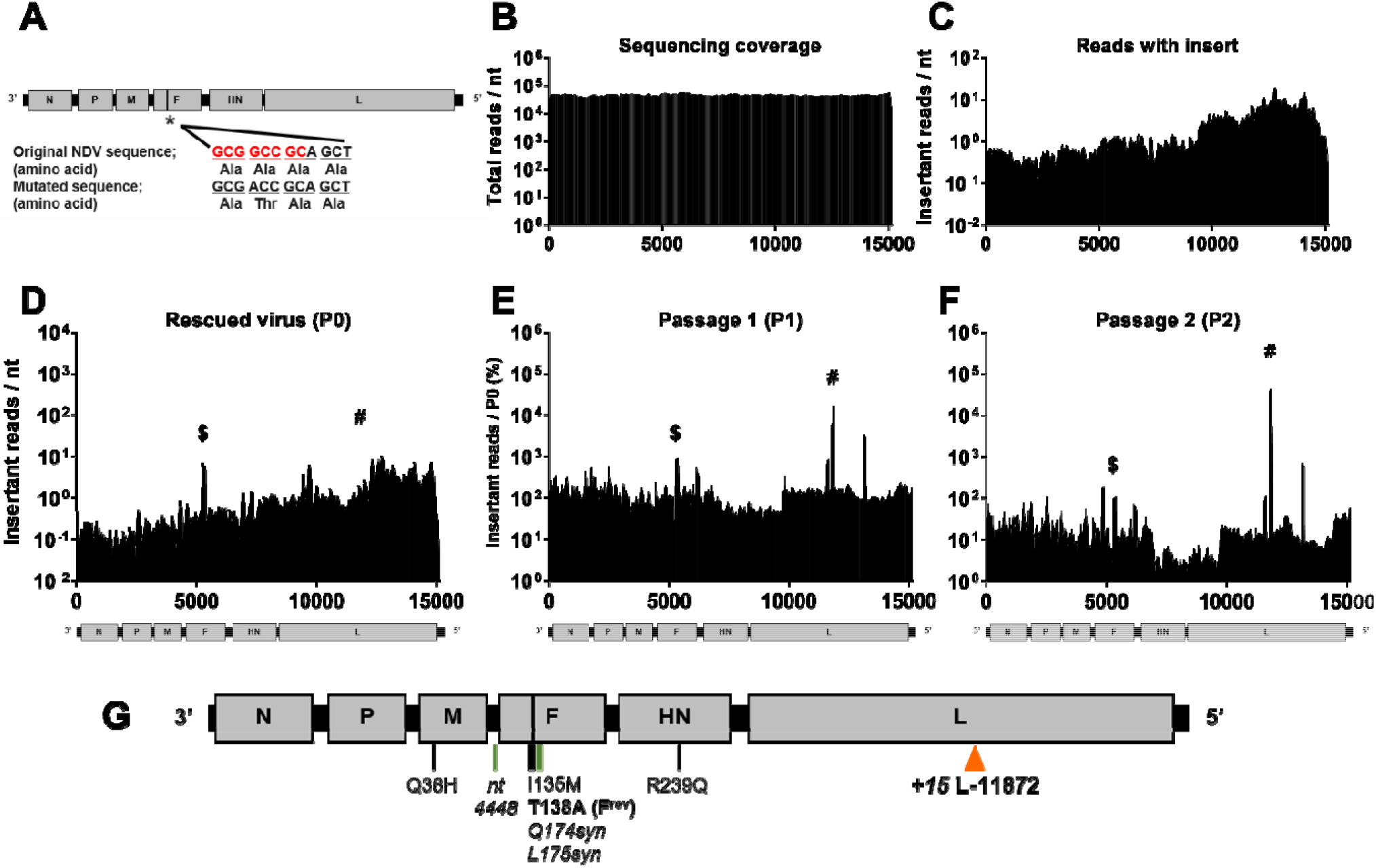
Insertional mutagenesis of fusion-defective Newcastle disease virus (NDV^Fmut^) selects for rare insertants in F and L genes stochastically associated with replicative fitness. **(A)** Elimination of the NotI restriction site (red sequence) in the NDV (6n+3) plasmid DNA with a single nucleotide change and concomitant mutation (*) Ala138Thr in the fusion peptide of F to generate NDV^Fmut^ (6n+3) for library generation. The vertical black bar in F indicates the location of the fusion peptide. **(B)** Total number of reads (y-axis) at each nucleotide position in the SeV genome, regardless of transposon detection, from the input plasmid. **(C-F)** Transposon insertion distribution within a 100nt sliding window, identified in (C) the NDV^Fmut^ plasmid DNA library input, (D) P0 (E) P1, and (F) P2. A to-scale schematic of the NDV genome is included under each graph defined as in Figs 2A-C and 4A-C. The y-axis in (C) and (D) represents the average number of reads containing an insertion (insertions per nt) at each nt position within a sliding 100-nt window surrounding the genome nt position on the x-axis. (E) P1 and (F) P2 y-axes, respectively, are normalized to the average read counts per nucleotide over three replicates expressed as percent of P0 input in (D) as was described for Figs 2B and 2C. Throughout (F and G), $indicates the insertants surrounding position 5383, and # indicates the insertants surrounding nts 11867-11877. **(G)** To-scale genome of the plaque-purified NDV-L-11872 clone. The fusion peptide is indicated by the black bar within F, as in (A) above. Transposon insertion at L-11872 is indicated by an orange arrow. Adaptive point mutations are indicated below the genome; green bars indicate mutations that do not affect amino acid sequence. T138A reverts the F^mut^ back to the wild-type, fusion-competent (F^rev^) sequence.

Our NDV^Fmut^ genomic plasmid library had a serendipitous skew in insertions towards the 5’ end of the genome that was not caused by sequencing bias (compare Fig. 6B to 6C). We also observed that this skew was maintained in the P0 rescue population (Fig. 6D), which reflects the likelihood that a wide range of NDV insertants were competent for genome amplification and budding. This suggests that NDV, like SeV, has a high overall capacity for genetic plasticity. However, the P0 infectious titer was extremely low at 10 iu/mL (Supplementary Table S4). This is expected since the vast majority of rescue events from the NDV^Fmut^ genomic library would result in the production of non-infectious virion particles. To further increase the selection pressure by genetic “bottlenecking”, NDV^Fmut^ (6n+18) P1 was carried out at an extremely low moi (<10^-5^). We observed a clear response to the bottleneck selective pressure upon subsequent passaging (P1 and P2, Fig. 6E and F), where the capacity for productive entry and fusion is essential for viral fitness, replication, and eventual amplification under the conditions examined.

### NDV^Fmut^ background selects for compensatory insertants in fusion protein

Remarkably, when we analyzed the insertant enrichment over two passages (Fig. 6D-F, Supplementary Table S5), we found that a vast majority of insertants were clustered around nts 11867-11877 in the L gene, with a subset of F insertants clustered around nt 5383 in NDV-F. This unusual distribution demonstrates that our experimental system is not inherently biased towards selection of insertants in the 3’ end of paramyxovirus genomes. As we had with our previous libraries, we recreated individual selected insertants and attempted rescue (Supplementary Table S5), but found that only the F-insertants were competent for virus spread while still maintaining the original F^A138T^ fusion-inactivating mutation in the NDV^Fmut^ genome (*e.g*. F-ORF-5367 and F-ORF-5384, Supplementary Table S5). These insertants correspond to the hinge region between domains III and I in the fusion protein, which has been implicated in fusion regulation (*17*), Thus, we identified insertants that specifically compensated for the alteration of the highly conserved A138 residue in the F-protein fusion peptide.

### Input transposon distribution drives selection of L-insertants associated with F^mut^-revertants

In addition to insertants that directly compensated for our fusion-peptide mutation, our hypothesis predicts that enrichment of other apparently viable insertants should occur on fusion-revertant genomes. Any such insertants should also follow the frequency distribution of the original input library. For example, since the transposon coverage of the input library was skewed towards the 5’ end of the NDV^Fmut^ genome by approximately ten-fold relative to the 3’ end of the genome (Fig. 6C), then any potential NDV-F^mut^-reversion and/or compensatory point mutations should also be more likely to occur in accordance with the probability distribution associated with 5’-skewed insertants. This is relevant when examining the highly-represented L-insertants. We noted that these insertants’ genomes were replication competent in the rescue cells, but did not produce infectious virus particles (Supplementary Table S5). When we re-examined the deep sequencing results *in toto* from NDV P2, we further identified high-abundance single nucleotide mutations in the NDV structural genes (M, F, and HN, the latter now formally designated as RBP (*2*)). Since L-ORF −11872 constituted such a high proportion of insertants in P2 (Fig. 6F), we double-plaque-purified NDV clones directly from P2 supernatant, and fully sequenced a genome containing the L-ORF-11872 insertant. Any such genomes in P2 are, by definition, viable and capable of spreading. Indeed, we found that in this clone, among other mutations in the structural genes, our original NDV^Fmut^ (F^A138T^) was reverted to the parental sequence, restoring the NotI sequence (Fig. 6G). While the significance of the other point mutations is unclear, the apparent fitness of L-ORF-11872 in the library pool is likely due to its co-occurrence on the same genome as the NDV^Fmut^ revertant.

While it is also conceivable that the L-insertant reduced the fidelity of the viral polymerase and permitted accumulation of compensatory mutations in the viral structural genes, several attempts to rescue this insertant alone (and others genetically nearby; Supplementary Table S5) in the NDV^Fmut^ background did not yield infectious virus, despite providing a wild-type L gene in *trans* during virus rescue to permit some replication and accumulation of compensatory mutations. In contrast, reversion of the NDV-F^mut^ to its wild-type counterpart (NDV-F^rev^) in the L-ORF-11872 insertant permitted virus rescue, amplification, and syncytia formation (Supplementary Table S5). Thus, reversion of the F^A138T^ point mutation was most likely responsible for the L-insertant’s relative fitness within the NDV library.

## Discussion

In order to explore the genetic plasticity of sialic acid-using paramyxoviruses, we generated saturating transposon insertional mutagenesis libraries of SeV and MuV, and then rescued and passaged these libraries to select for relatively fit insertants. We found that both SeV (Fig. 2) and MuV (Fig. 4) tolerated insertions in the N and P genes, and especially in their untranslated regions. When we rescued selected clones of the most highly-enriched insertants, we found that overall capacity for rescue correlated with their original enrichment in the library (Figs. 3 and 5), indicating that the screening libraries successfully predicted insertant viability and identified broadly-plastic genetic regions. However, in virus growth and competition assays, both the SeV and MuV insertants showed differential fitness in comparison to their representation in the original screening library (Figs. 3G, 5G), indicating that abundance within the screening library only poorly predicts relative fitness of individual genomes. With these libraries, overall we are able to draw widely-applicable conclusions about the broad genetic plasticity of paramyxoviruses. In addition, by employing a separate Newcastle disease virus library where we introduced a selective pressure to bias insertant distribution differently than what was observed with SeV and MuV, we demonstrate the potential of these libraries for genome-wide interrogation of paramyxovirus fitness landscape.

Broadly, the data from these libraries correlate well with our earlier work on insertional mutagenesis of MeV. Our original intent in adding SeV and MuV libraries to our insertional mutagenesis repertoire was to identify whether usage of sialic acid would permit greater tolerance to structural change (*11*) or whether we would still observe significant constraint on PMV glycoproteins that could explain their well-known lack of antigenic drift (*3*) as we saw with MeV (*10*). We found that SeV and MuV do not show increased tolerance for insertions in their glycoproteins. This is in contrast to the genetic and structural plasticity observed in the HA (hemagglutinin) glycoproteins of sialic acid-using Influenza viruses (*11, 18*), indicating that receptor usage does not determine tolerance to insertional mutagenesis. Even when we disabled NDV fusion as a means of forcing change in the virus’ structural region, we observed a strong selection for reversion mutants, and only a lesser accumulation of compensatory insertants. When we tested the viability of individual insertants in these regions with SeV and MuV, we found that these viruses were incompetent for virus amplification. These observations are consistent with the monoserotypic nature of most PMV species (*3–6, 8*), and with careful analysis of evolutionary constraints on PMV fusion proteins (*19*). Despite using overlapping receptors, orthomyxoviruses (including influenza viruses) and paramyxoviruses have significantly different entry strategies; while the influenza virus HA glycoprotein coordinates both receptor binding and fusion in a single protein, it requires a separate protein (NA, neuraminidase) to release virions from infected cells. Paramyxoviruses, in contrast, contain neuraminidase (when needed) and receptor binding activities in a single protein (RBD), while using a separate protein (F) to initiate membrane fusion, and there is a tightly co-ordinated series of interactions between the RBD and F proteins during virus entry that likely varies among different paramyxovirus genera (*20*). This co-ordinated interaction process likely introduces stringent constraints on both proteins to preserve interactions and capacity for structural rearrangements that are not present in influenza virus glycoproteins. This further confirms that there are broad structural constraints on PMV glycoproteins that prevent them from undergoing the antigenic drift observed in orthomyxoviruses like Influenza A virus (*9*).

All three viruses have high insertant coverage in the input plasmid library, but only SeV and NDV P0 insertant coverage remained high upon initial rescue, while MuV P0 shows a stark drop-off in total insertants and a shift in insertant distribution from the input. This is likely a representation of the underlying genetic plasticity of each genome, rather than an effect of rescue efficiency, based on data from another genome library screen with Nipah virus (NiV) as a representative of the genus *Henipavirus* (not shown). Rescue and amplification of NiV transposon mutagenesis libraries was carried out under BSL-4 conditions, precluding optimization of rescue efficiency to the same degree as we had done with the other PMVs. And so, while input plasmid insertant coverage was similar in depth and breadth to SeV, MuV, and NDV, only ~5500 rescue events occurred. Insertant distribution was stochastic due to this low efficiency, but insertants were detected broadly throughout the genome, demonstrating that poor rescue efficiency does not drive shifts in insertant distribution. Thus, the change in MuV insertant distribution from input to P0 is likely representative of biological restrictions on whether the genomes could be rescued. SeV and NDV do not share this level of restriction, indicating that these viruses’ genomes are more plastic.

Insertant distribution after passaging in the fusion-competent SeV, MuV, and MeV genomes demonstrate two related patterns of enrichment: firstly, more viable insertants are located towards the 3’ end of these genomes (Figs 2A-C, 4A-C), and secondly, the viruses are generally more enriched for insertants in the UTRs of the genes than their cognate ORFs, especially in transcriptional units like N and P, where there is sufficient insertional coverage at P2 to make such comparisons (Figs 2E and 4E). The observation of increased insertional tolerance in UTRs is not surprising; PMV UTRs play a regulatory role in transcription, mRNA stability, and translation efficiency, in ways that are not thoroughly characterized, but they are overall only constrained at the nucleotide level. However, even within this, there is still a clear and overriding 3’ enrichment bias, since HN and L UTRs of SeV, MuV, or MeV do not demonstrate insertant enrichment. Even the NDV^Fmut^ library does not demonstrate an enrichment in these UTRs, despite the library’s input bias and ultimate enrichment for insertants in a presumed “neutral” region of the adjacent L-ORF. We therefore hypothesize that highly-expressed genes like N and P can tolerate some dysregulation without significant negative effects, whereas the intolerance to dysregulation of less-abundant genes like F, H/HN, and L may be an indicator of how stringently-regulated they are. Our studies suggest that the regulatory functions of these intergenic regions in PMVs (*21–24*) should be systematically explored in their appropriate genomic contexts, which is now possible using robust and efficient reverse genetics systems.

In SeV and MuV, these enriched UTRs near the 3’ end of the genome are co-incidentally near the eGFP reporter gene, which is between N and P. Nonetheless, this likely reflects an aspect of virus biology rather than a simple proximity to the reporter gene since we also previously observed an increased tolerance for insertants in the 3’ end of the MeV genome (*10*). In addition, the eGFP reporter in our NDV reporter genome is located between P and M, but we do not see enrichment for insertion in those untranslated regions.

We also observed an enrichment for insertions in the coding regions of N and P in our fusion-competent libraries. Paramyxoviral P proteins code for multiple accessory proteins in different frames by use of alternative translation start sites (C proteins) and by mRNA editing (V, W), and these proteins are largely involved in blocking host antiviral sensing and response. Despite the constraint of coding in multiple frames, we observed enrichment for insertions in all our fusion-competent libraries at the 5’ (N-terminus) end of the P gene, the region common to all the ORFs. However, the C, V, and W proteins of PMVs bear unstructured regions and are highly variable between virus species and genera (*1, 25, 26*), while the N-terminus of P is specifically understood to be intrinsically disordered (*27*). Together, this may explain the unexpected insertional tolerance in P. N insertions are also found primarily in the unstructured C-terminus “Ntail” region, which has already been shown to accommodate insertions and deletions with limited negative impact on MeV in tissue culture (*28, 29*). We thus propose that our transposon mutagenesis enriches for insertants in such unstructured regions of proteins, since specific functional elements will remain accessible regardless of upstream and downstream insertions.

The NDV^Fmut^ library P2 insertants indicate that structural order, however, is not the only determinant of insertion tolerance in coding regions. Dochow *et al*. produced an analysis of the propensity for disorder across the MeV L protein as a model for other PMVs, and further tested select predicted unstructured regions for tolerance to small insertions and epitope tags (*30*). Although the majority of the NDV L-ORF was enriched for transposon insertions in the NDV^Fmut^ DNA input library, over passage only a small subset of closely-located L-insertants were viable when combined with revertant F point mutations. Negative-sense RNA virus polymerases contain six major conserved regions (CRs) flanked by variable regions that differ between and within virus families. The NDV^Fmut^ L-insertants are all within a small portion of CR-IV, the function of which is not understood. Based on Dochow *et a’s* analysis, this region is generally ordered, and so it is not obvious why insertants in this region are more viable. Other nearby portions of L are predicted to be far more disordered, and in fact an unstructured region near CR-VI has been shown to tolerate both large epitope tags and a complete break of the polymerase into two separate ORFs, as long as they are brought back into contact by artificial domains (*30*). While we cannot determine causation within the library setting, follow-up failed attempts to rescue the L-insertants alone without revertant F mutations does not suggest that these insertants specifically potentiated acquisition of point mutations; *i.e*. we can find no evidence that the enriched L-insertants are more error-prone polymerases. Thus, a much more detailed structural analysis and mutagenesis exploration of NDV-L, as well as other PMV polymerases, will be required to understand what determines this region’s specific tolerance to insertion. It is interesting to note that since this region was not predicted by structural analyses, a functional insertional mutagenesis assay does still have information to offer for designing sites for tagging viral proteins or inserting novel tandem ORFs and fusion proteins.

Within this body of work, we compared fitness in the screening libraries with individual insertant clonal fitness, and competitive fitness in the more-contained competitive outgrowth assays. We found that relative clonal fitness (as measured by growth curves and peak titer) correlated well with relative fitness in competitive outgrowth assays, particularly for SeV. However, in the context of the library setting, the number of insertant reads at P2 appears to be more affected by complex epistatic factors. The ranked frequency of insertant reads at P2 did not always match their clonal or competitive fitness in more controlled assays. Altogether, the evidence suggests that the transposon library approach, without the more careful downstream analyses shown in our studies, is best suited to dissecting viable *vs* non-viable viral genomes in our assay setting, rather than predicting the relative fitness of individual insertants.

By placing our NDV^Fmut^ library under a unique form of selective pressure, we drove enrichment of insertants in otherwise-intolerant regions of the genome – F, a structural protein, and L, the viral polymerase. Thus, we have demonstrated the power of our strategy to reveal not only the genomic plasticity of paramyxovirus genomes, but also the ability to use our methods to design arbitrarily selective screening campaigns to interrogate paramyxovirus biology. Specifically, we envision leveraging our efficient and robust reverse genetics systems to design and execute selection strategies that can be used to interrogate the fitness landscapes of individual genes that were previously not accessible using conventional paramyxovirus passaging and selection. Furthermore, we have shown the viability of designing strategies to select for mutants in fitness landscapes that are otherwise not easily accessible during the normal course of paramyxovirus evolution.

*In toto*, we found common regions of tolerance and intolerance for insertions in PMV genomes, specifically identifying tolerance to dysregulation of highly-expressed genes. We further noted that there are structural constraints on changing PMV antigenicity, and that unstructured regions in the N, P, and accessory proteins permit insertional mutagenesis. We demonstrated that this highly-disruptive whole-genome insertional mutagenesis library approach could be informative for paramyxoviruses placed under unique selective pressures: not only such genetic pressures as we used here, but also perturbations like interferon treatment or amplification in susceptible host animals. Overall, the combined commonalities and differences in these paramyxovirus mutagenesis libraries provide a broader family-wide context in which to understand specific variations within PMV genera or species.

## Materials and Methods

### Experimental Design

Our transposon mutagenesis strategy for whole genome interrogation of paramyxoviruses is outlined in Fig. 1 and the accompanying text. It takes advantage of our efficient and robust reverse genetics system (*31*) and leverages the rule-of-six (*15*), the latter being a unique feature of paramyxovirus replication. Whole genome insertional mutagenesis libraries were generated for three paramyxoviruses described below, and these libraries were rescued (P0) and passaged (P1, P2) in tissue culture to identify genetic regions that were relatively tolerant to insertion for downstream characterization.

#### Research Objectives

Through this study we sought (i) to identify genetic regions or determinants of plasticity common to sialic-using paramyxoviruses, (ii) to explore how determinants of fitness within library settings correspond to solo and alternative library selective and competitive pressures, and (iii) to test the effects of a defined selective pressure on which genetic regions would demonstrate plasticity.

(i) Genome-wide transposon mutagenesis screens have been carried out on other positive and negative sense RNA viruses, including our own previous study on MeV with similar goals (*10*).

(ii) Validation of such an approach usually includes only generation of recombinant viruses bearing select enriched insertants to ensure that they are viable. However, in the context of non-segmented negative sense RNA viruses such as PMVs, the replicative fitness of individual insertants derived from a complex pool may not be accurately reflected in their individual growth kinetics and peak titers. To gain a better understanding how the abundance of the insertants identified in the library P2 relate to growth kinetics and replicative fitness, we made recombinant 6n+18 PMVs bearing representative insertants from the relevant genomic regions. We then determined their individual growth kinetics and peak titers, and subjected them to a competitive outgrowth assay. We developed a Nanopore long-read sequencing protocol and bioinformatic pipeline to monitor the outcome of a focused multiplex competitive outgrowth assay. This competitive outgrowth assay revealed substantive differences between how the abundance of the SeV or MuV insertants detected in library P2 Illumina reads relate to their solo replicative fitness and competitive fitness in a more focused assay.

(iii) Finally, to determine that our whole genome transposon mutagenesis screens of paramyxoviruses was not systematically biased towards detecting only 3’ end genomic insertants in the N-P gene regions, we applied our screen to a parental 6n+3 NDV genome that was made fusion-defective (NDV^Fmut^) by destroying a naturally occurring NotI site in the genetic region encoding the fusion peptide in NDV-F. To make a *Mu*-transposon mutagenesis library, the parental (6n+3) genomic plasmid has to be devoid of NotI sites. Subjecting the NDV^Fmut^ transposon mutagenesis genomic library to rescue and passaging resulted in identification of compensatory F-insertants in an otherwise fusion-defective background, along with a cluster of L-insertants that occurred on a F-revertant fusion-competent background. These results show the power of experimental system to design arbitrarily selective screening campaigns and access distant fitness landscapes to interrogate paramyxovirus biology.

#### Units of investigation

##### Viruses

We selected SeV^Fushimi^, MuV^JL5^, and NDV^LaSota^ for use in this study as sialic acid (SA)-using representative viruses of major paramyxovirus subfamilies *Orthoparamyxovirinae, Rubulavirinae*, and *Avulavirinae*, respectively. The individual strains were selected based on previous optimization of high-efficiency reverse genetics systems for each virus (*31*).

##### Cells

We rescued our original transposon mutagenesis libraries in BSRT7 cells (derived from BHK21 cells) since we have achieved maximal rescue efficiency with our reverse genetics systems on these cells (*31*). We passaged the rescued output (P0) from SeV and MuV in Vero cells for two sequential passages (P1 and P2). Our NDV library was both rescued and amplified in BSRT7 cells since we observed better growth in these cells than in Vero cells. We reasoned that both BSRT7 and Vero cell lines, which are respectively deficient in interferon signaling (*32*) and interferon production (*33*), should serve as neutral cellular backgrounds for our initial genome-wide interrogations. Using these cells removes likely confounding selective pressures against insertions in the P gene that might disrupt its interferon antagonist function (and that of P-derived accessory proteins). We did not want to miss insertants in the P gene that might be structurally tolerated and not affect P’s function as a cofactor of the L-mediated transcriptase and replicase complex.

#### Sample size/scale

The rescue efficiency of our reverse genetics system was critical for determining which paramyxovirus could be interrogated on a genome-wide scale. If transposon insertions were truly random and not affected by other confounding factors (see footnotes to Supplementary Table S1), Poisson statistics dictate that a 10X coverage in terms of the number of independent transposon insertants rescued, relative to the size of the cognate PMV genome, is required to ensure >90% probability that any given nucleotide position in the genome has at least one insertant, so rescue scale was dictated by each virus’ rescue efficiency as calculated in Supplementary Tables S1 and S4.

#### Replicates

Library generation, and library and insertant rescue was carried out in singlicate as an appropriate use of resources. Insertants whose rescue failed to produce infectious viruses, two additional rescue attempts were made to verify the insertants’ inviability. All other assays were carried out in triplicate unless otherwise noted to ensure accurate representation of the potential diversity of assay outcomes.

#### Endpoint

In all library scenarios and in the multiplex competitive outgrowth assays, the endpoint for each passage was defined as when the entire culture was infected, as determined by visual observation of GFP-positivity. For individual insertant rescue, success or failure was determined by whether virus demonstrated spread in tissue culture within ten days of transfection – monitored by GFP-positivity, and confirmed by syncytia formation for MuV and NDV. Growth curves were carried out until the wild-type (6n) virus titers plateaued and began to reduce in magnitude.

#### Inclusion/Exclusion criteria

No samples or replicates were discarded from these experiments. Computational criteria for identifying insertants in deep sequencing results are based on the sequencing fidelity for the platform used (Illumina and Nanopore), and are otherwise as inclusive as possible to identify all insertants.

### Cell lines

Vero cells (ATCC Cat# CCL-81, RRID:CVCL_0059), and BSR T7/5 cells (RRID:CVCL_RW96; (*34*)) were propagated in Dulbecco’s modified Eagle’s medium (ThermoFisher Scientific, USA) supplemented with 10% fetal bovine serum (Atlanta Biologicals, USA) at 37°C. Cell lines were monitored monthly to maintain mycoplasma-negative status using the MycoAlert Mycoplasma Detection Kit (Lonza, USA).

### Plasmids and viruses

Genome coding plasmids for SeV (pSeV^Fushimi^-eGFP; KY295909), MuV (pMuV^JL5^-eGFP; KY295913), and NDV (pNDV^LaSota^-eGFP; KY295917) were modified to have optimized T7 promoter and hammerhead ribozyme as previously reported (*31*). Our recombinant SeV^Fushimi^-eGFP bears mutations in the M and F genes that enable trypsin independent growth (*35*). The MuV^JL5^-EGFP strain is derived from the JL5 vaccine strain. NDV-eGFP was based on LaSota strain with mutations in its cleavage site to be cleaved by urokinase-type plasminogen activator (uPA) (*36*). In order to attenuate viral genomes that lack the transposon insertion, we introduced an extra 3nt stop codon after the reporter gene in each construct, rendering the viral genome 6n+3 nucleotides; these are indicated as SeV 6n+3, MuV 6n+3, and NDV 6n+3. We also abolished NotI restriction sites in each virus’ plasmid in order to facilitate transposon removal. All modification for plasmids were performed using overlap PCR mutagenesis with InFusion cloning (Takara Biosciences, USA). Viral genome and support plasmids were maintained in chemically-competent Stbl2 *E. coli* cells (ThermoFisher Scientific, USA) with growth at 30°C.

Supplementary Table S6 contains the primer sequences for generating all recombinant insertant plasmid genomes. Nucleotide position in the genome is labelled without the eGFP transcriptional unit, and insertant position refers to the nucleotide after which the transposon sequence began. Insertants are named by genetic region and nucleotide position.

### Transposon-mediated mutagenesis

The Mutation Generation System (ThermoFisher Scientific, USA) was used to randomly insert transposons in the 6n+3 genomic plasmids using a modified protocol. An *in vitro* transposon insertion reaction was performed on approximately 850ng per viral genome plasmid (40ng DNA per kb of plasmid) of 6n+3 genomic plasmids, which were dialyzed twice for 30 min in 1L ddH_2_O, and then transformed into ElectroSHOX cells (Bioline USA, discontinued). Following transformation, the cells were plated on 20 x 15cm plates with LB agar containing ampicillin (MilliporeSigma, USA) and kanamycin (ThermoFisher Scientific, USA) (selecting for plasmid transformants and transposon insertion respectively) and allowed to grow for ~18 hours at 30°C. The bacterial colonies were scraped from the agar with PBS, and pelleted, and DNA was extracted from the pooled colonies using a PureLink HiPure maxiprep kit (ThermoFisher Scientific, USA). 30ug of transposon-containing genomic plasmid was digested with NotI-HF (New England Biolabs, USA) for 3 hours to remove the transposon body. The restricted plasmid was then gel purified using the Qiaex II kit (Qiagen, USA), and 500 ng of the DNA was re-ligated at 25°C for 30 minutes using T4 DNA Ligase (New England Biolabs, USA) and heat-inactivated at 65°C for 10 minutes. The entire ligation mixture was dialyzed for 20 min in 1L ddH2O, and then transformed into ElectroSHOX cells and plated on 20 x 15cm plates containing ampicillin only. After ~18 hours’ growth at 30°C, the colonies containing 6n+3 viral genomes with the transposon scar (6n+18) were again scraped from the plates into PBS, and the viral 6n+18 genome DNA was again extracted using the HiPure maxiprep kit.

### Rescue of recombinant viruses (P0) from cDNA

For recovery of recombinant viruses, rescue was performed as described in Beaty et al. (2017). 4×10^5^ BSR T7/5 cells per well were seeded in 6-well plates. The following day, DNA and Lipofectamine LTX / PLUS reagent (ThermoFisher Scientific, USA) were combined as indicated in Supplementary Table S7 in OptiMEM (ThermoFisher Scientific, USA) with gentle mixing by pipetting only. After incubation at room temperature for 30 minutes, the DNA:lipofectamine mixture was added dropwise onto cells. Separate transfection reactions were set up for each rescue well. Transfected cells were incubated at 37°C for 8-10 days, until the level of infection reached 100% as determined by observation of GFP-positive cells by microscopy. Supernatant was collected from rescue cells, pooled, and clarified by centrifugation. Clarified supernatants were stored at −80°C.

### Analysis of relative rescue efficiency

SeV-WT, SeV-parental (6n+3), and SeV-library (6n+18) genomes were rescued as described in detail above. Two days post-rescue, cells were collected with PBS+50mM EDTA, pelleted, and re-suspended into 2% paraformaldehyde for fixation. After 15 minutes, cells were pelleted again, and re-suspended into PBS + 2mM EDTA + 2%FBS. Cells were assayed by flow cytometry on a BD FACSCantoII with BD FACSDiva v6.0, and evaluated for GFP-positivity in the Blue-1 channel, relative to un-transfected cells. 5×10^5^ events were collected for each sample - the equivalent of a full 6-well well. The WT, parental (6n+3), and libraries (6n+18) for MuV were evaluated the same way.

### Titering viral supernatants

Titrations of SeV, MuV, and NDV stocks were performed on Vero cells in a 96-well format, with individual infection events (infectious units, iu/mL) identified by GFP fluorescence at 24 hours post-infection using an Acumen plate reader (TTP Labtech, USA).

### Passaging virus for SeV and MuV library screen

5.2×10^6^ Vero cells in a 15cm dish were infected at an MOI of 0.01 for each passage and replicate, with the exception of passage 1 in MuV. We adopted a MOI of 0.0003 (5120 iu/dish) for passage 1 (P1) of MuV, because P0 titer was very low. Thereafter, infection was monitored by microscopy and supernatants were collected when the level of infection reached 100% as determined by observation of GFP-positive infected cells by microscopy, at 8-10 days post-infection (dpi).) Supernatant was collected from rescue cells, pooled, and clarified by centrifugation. Clarified supernatants were stored at −80°C. Screening experiments were done in triplicate independently.

### Passaging virus for NDV library screen

BSRT7 cells were infected with NDV using the same strategy and parameters as with SeV and MuV above. Since infectious titers from P0 (rescue) of NDV were very low, P1 was carried out at a very low moi (<0.0001) and required several additional days to reach confluence post-infection.

### RT-PCR and Illumina sequencing for library screen

The SeV, MuV, and NDV RNA was extracted from thawed supernatant using QIAamp viral RNA extraction kit (Qiagen, USA). Genomic RNA was then amplified in six equal-sized segments using overlapping primers sets (Supplementary Table S8) using SuperScript III One-Step RT-PCR kit (ThermoFisher Scientific, USA) with Platinum Taq. The cDNA segments from each sample were pooled in equimolar amounts, sheared by Covaris sonication, and prepped for sequencing using TruSeq DNA LT Sample Prep Kit (Illumina, USA) according to the manufacturer’s instructions. Barcoded and multiplexed samples were sequenced on a HiSeq2000 using 100-nt single-end reads in Rapid Run mode. Analysis of the transposon insertions was performed as previously described (*10*).

### Sequencing analysis of library screen

Identification of the transposon insertions were carried out as in Heaton *et al. (11*). Briefly, reads with the transposon scar sequence of TGCGGCCGCA were extracted from the total sequencing data. The scar sequence was then deleted, leaving a 5nt duplication at the site of insertion. These sequences were then aligned to the viral reference sequences by bowtie2, and processed sam files were used to identify the position of insertion in each read.

### Data analysis and insertant selection from library screens

Although transposon coverage was overall >10-fold for both SeV and MuV libraries, individual nucleotide positions did not always receive an insertion. Thus, all analyses were carried out using a 100nt sliding window to prevent division by 0. Additionally, raw insertant counts in P1 and P2 are likely to be biased by varying transposon abundance in the input and rescue (P0) libraries, so we normalized P1 and P2 reads by the number of insertants in P0, and presented these passages as triplicate average percent reads over P0 (Figs. 2A-C and 4A-C).

To identify the most highly-enriched individual insertants in the library, we first identified 40 insertants with the highest overall raw read count at P2 from each library. We then divided these by normalized P0 reads, and eliminated any insertants whose relative abundance drastically decreased over passage (average P2/P0 < 30%) in order to account for variability of coverage in P0. From those remaining, we showed the top 20 insertants for SeV (Supplementary Table S2), and the insertants that showed an average of 1 or more reads in MuV (Supplementary Table S3). Individual insertants from these lists were selected for downstream characterization as described in Results.

### Insertant rescue and growth curves

Individual insertant viruses were rescued in BSR T7/5 cells as described above, were amplified in Vero cells once, and titered as above. 2×10^5^ Vero cells per well in a 12-well dish were infected at an MOI of 0.01 for 2h, followed by replacement of fresh medium. Samples were collected daily for titration with complete media exchange.

### Competitive outgrowth assay

Because individual insertant viruses demonstrated different growth characteristics that could render our standard titration assay (described above) inaccurate, we titered the individual insertants by focus-forming assay prior to combining them for a competition outgrowth assay. 2×10^5^ Vero cells per well in 12-well plates were inoculated with a serial 10-fold dilution of insertants for 2 hours. Cells were washed with PBS once and then replaced with an overlay methylcellulose (1% methylcellulose in DMEM plus 2% FBS) to prevent establishment of secondary foci. At 7 dpi (SeV) or 4 dpi (MuV), the number of eGFP-positive infectious foci were manually counted using a Nikon Eclipse TE300 inverted fluorescent microscope (Melville, NY, USA).

Competitive outgrowth assays were carried out in independent biological triplicates: equal infectious units as defined by the focus-forming assay above of 8 SeV insertants or 6 MuV insertants were mixed, creating P0 mixture. Then the titer of each of these mixtures was re-quantified by iu as described above. 2×10^6^Vero cells in a 10 cm dish were infected by the P0 mixture at a final MOI of 0.01. In order to maintain cell viability, media was exchanged daily until ~100% infection was reached as determined by eGFP-positive cells by fluorescent microscopy. The supernatant on the day of eGFP confluency was used as the P1 sample. For P2, supernatant from P1 was titered (by infectious unit), then inoculated at MOI of 0.01 and passaged until 100% infection as was done for P1.

### RT-PCR and Nanopore sequencing for competitive outgrowth assay

RNA was extracted from P0, P1, and P2 viral supernatant and the relevant genetic regions encompassing all the insertants for a given virus were reverse-transcribed and amplified as described for Illumina sequencing. These amplicons were prepared for Nanopore sequencing using the native barcoding expansion kit (EXP-NBD104, Oxford Nanopore Technology, United Kingdom) and ligation sequencing kit (SQK-LSK109, Oxford Nanopore Technology, United Kingdom), and then sequenced on a MinION R9.4.1 flow cell (Oxford Nanopore Technology, United Kingdom) to determine the abundance of insertants in each sample and passage.

### Sequencing analysis of competitive outgrowth assay

Nanopore basecalling was carried out using Albacore, then aligned to reference sequence by Burrows-Wheeler Aligner. Since MinION DNA sequencing is prone to error, we identified the insertants by extracting inserts ≥ 10nt in size, and extracted the number of insertions at each position. Abundance of each insertant was calculated relative to the total insertion count.

### Correlation analyses

Insertants represented in all three assays (library screen, peak titer, and competitive outgrowth) were analyzed for their relative fitness across assays. Since each assay used metrics with different ranges of magnitude, comparison across assays was facilitated by ranking insertants within each assay (see below) resulting in non-parametric distributions. Assays were thus compared by Speaman’s correlation analysis and r values as measures of correlation are reported. * P <0.05.

*Ranking: (a) Library screen:* Insertants were ranked according to their raw abundance at P2 in each replicate, and then the average of the three replicates were used to define the insertants’ rank. (*b) Peak titer:* Insertants were ranked by peak titer in each growth curve repeat, and the average of the three repliactes were used to define the insertants’ rank. (*c) Competitive outgrowth:* Insertants were ranked by magnitude of expansion (input/P2) in each replicate, and the average of the three replicates were used to define the insertants’ rank.

## Supplementary Materials

**Supplementary Table S1.** Transposon mutagenesis calculations and metrics of SeV and MuV.

**Supplementary Figure S1.** Sequencing and transposon coverage of SeV library.

**Supplementary Table S2.** Most highly-represented insertants from SeV library.

**Supplementary Figure S2.** Sequencing and transposon coverage of MuV library.

**Supplementary Table S3.** Most highly-represented insertants from MuV library.

**Supplementary Table S4.** Transposon mutagenesis calculations and library metrics in NDV.

**Supplementary Table S5.** Most highly-represented insertants from NDV library.

**Supplementary Table S6.** Primers for generation of insertant clones.

**Supplementary Table S7.** SeV, MuV, and NDV rescue parameters.

**Supplementary Table S8.** Primers for one-step reverse-transcription and PCR of SeV, MuV, and NDV.

## General

We thank Dr. W. Paul Duprex and Dr. Nancy McQueen for the kind gift of the MuV and SeV reverse genetics systems we previously modified to make this manuscript possible. Both were co-authors in the original paper describing the efficient and robust reverse genetics system that we developed (*31*). We also thank other members of the Lee Lab for their helpful feedback, and Dr. Benjamin tenOever for help with some of the original stream graphs. We would also like to acknowledge the genomics and flow cytometry cores at the Icahn School of Medicine at Mount Sinai.

## Author Contributions

S.M.B. and B.L. conceived the experiments. S.M.B., P.A.T., A.P., S.T.W., and P.H. generated insertional libraries and carried out screening. S.M.B carried out library rescue and passaging. S.I. performed the critical analyses, generated individual virus clones based on these analyses, and carried out competition assays. S.I. and B.L. conceived and analyzed, but S.I. executed the NDV validation screens. D.S. and C.S. aided in sequencing data analysis. C.S. wrote the code to identify the inserts from the MinION platform (Oxford Nanopore). P.A.T., S.I., and B.L. wrote the manuscript.

## Data and materials availability

The raw next generation sequencing results are uploaded at NCBI GEO. Library screening Illumina sequencing: GSE125257 for SeV and MuV, GSE138366 for NDV. Competitive outgrowth assay Oxford Nanopore sequencing: GSE128625. SeV-eGFP and NDV-eGFP may be obtained through a MTA with Icahn School of Medicine at Mount Sinai via Mount Sinai Innovation Partners (MSIP). MuV (JL5)-eGFP needs to be requested first through Dr. W. Paul Duprex, Center for Vaccine Research, University of Pittsburg, then through the corresponding author (Dr. Benhur Lee).

## References

1. P. A. Thibault, R. E. Watkinson, A. Moreira-Soto, J. F. Drexler, B. Lee, in Advances in Virus Research. (2017), vol. 98.

2. B. Rima et al., ICTV Virus Taxonomy Profile: Paramyxoviridae. Journal of General Virology 100, 1593–1594 (2019).

3. S. M. Beaty, B. Lee, Constraints on the Genetic and Antigenic Variability of Measles Virus. Viruses 8, 109 (2016).

4. M. Tsurudome, M. Nishio, H. Komada, H. Bando, Y. Ito, Extensive antigenic diversity among human parainfluenza type 2 virus isolates and immunological relationships among paramyxoviruses revealed by monoclonal antibodies. Virology 171, 38–48 (1989).

5. K. L. van Wyke Coelingh, C. C. Winter, E. L. Tierney, W. T. London, B. R. Murphy, Attenuation of bovine parainfluenza virus type 3 in nonhuman primates and its ability to confer immunity to human parainfluenza virus type 3 challenge. J Infect Dis 157, 655–662 (1988).

6. H. Q. McLean, A. Parker Fiebelkorn, J. L. Temte, G. S. Wallace, Prevention of Measles, Rubella, Congenital Rubella Syndrome, and Mumps, 2013. MMWR Recommendations & Reports 62, 1–35 (2013).

7. J. F. Drexler et al., Bats host major mammalian paramyxoviruses. Nature Communications 3, 796 (2012).

8. S. M. Beaty et al., Cross-reactive and cross-neutralizing activity of human mumps antibodies against a novel mumps virus from bats. Journal of Infectious Diseases pii: jiw53, (2016).

9. M. O. Altman, D. Angeletti, J. W. Yewdell, Antibody Immunodominance: The Key to Understanding Influenza Virus Antigenic Drift. Viral Immunol 31, 142–149 (2018).

10. Benjamin O. Fulton et al., Mutational Analysis of Measles Virus Suggests Constraints on Antigenic Variation of the Glycoproteins. Cell Reports 11, 1331–1338 (2015).

11. N. S. Heaton, D. Sachs, C.-J. Chen, R. Hai, P. Palese, Genome-wide mutagenesis of influenza virus reveals unique plasticity of the hemagglutinin and NS1 proteins. Proceedings of the National Academy of Sciences 110, 20248–20253 (2013).

12. B. K. Rima et al., Stability of the Parainfluenza Virus 5 Genome Revealed by Deep Sequencing of Strains Isolated from Different Hosts and following Passage in Cell Culture. Journal of Virology 88, 3826–3836 (2014).

13. M. Mateo, C. K. Navaratnarajah, R. Cattaneo, Structural basis of efficient contagion: measles variations on a theme by parainfluenza viruses. Curr Opin Virol 5, 16–23 (2014).

14. P. Plattet, R. K. Plemper, Envelope Protein Dynamics in Paramyxovirus Entry. mBio 4, e00413–00413 (2013).

15. P. Calain, L. Roux, The rule of six, a basic feature for efficient replication of Sendai virus defective interfering RNA. Journal of Virology 67, 4822–4830 (1993).

16. Y. Shirogane, S. Watanabe, Y. Yanagi, Cooperation between different RNA virus genomes produces a new phenotype. Nature Communications 3, 1235 (2012).

17. M. Chi et al., Conserved amino acids around the DIII-DI linker region of the Newcastle disease virus fusion protein are critical for protein folding and fusion activity. BioScience Trends 13, 225–233 (2019).

18. B. O. Fulton, W. Sun, N. S. Heaton, P. Palese, The Influenza B Virus Hemagglutinin Head Domain Is Less Tolerant to Transposon Mutagenesis than That of the Influenza A Virus. J Virol 92, (2018).

19. V. A. Avanzato et al., A structural basis for antibody-mediated neutralization of Nipah virus reveals a site of vulnerability at the fusion glycoprotein apex. Proceedings of the National Academy of Sciences 116, 25057–25067 (2019).

20. K. D. Azarm, B. Lee, Differential Features of Fusion Activation within the Paramyxoviridae. Viruses 12, (2020).

21. J. C. Rassa, G. M. Wilson, G. A. Brewer, G. D. Parks, Spacing Constraints on Reinitiation of Paramyxovirus Transcription: The Gene End U Tract Acts as a Spacer to Separate Gene End from Gene Start Sites. Virology 274, 438–449 (2000).

22. A. Sugai, H. Sato, M. Yoneda, C. Kai, Gene end-like sequences within the 3’ non-coding region of the Nipah virus genome attenuate viral gene transcription. Virology 508, 36–44 (2017).

23. K. Hino et al., Downregulation of Nipah virus N mRNA occurs through interaction between its 3’ untranslated region and hnRNP D. Journal of virology 87, 6582–6588 (2013).

24. Y. Inoue et al., Selective Translation of the Measles Virus Nucleocapsid mRNA by La Protein. Frontiers in Microbiology 2, (2011).

25. M. K. Lo, T. M. Søgaard, D. G. Karlin, Evolution and Structural Organization of the C Proteins of Paramyxovirinae. PLoS ONE 9, e90003 (2014).

26. Y. Fujii, K. Kiyotani, T. Yoshida, T. Sakaguchi, Conserved and non-conserved regions in the Sendai virus genome: evolution of a gene possessing overlapping reading frames. Virus genes 22, 47–52 (2001).

27. J. Habchi, S. Longhi, Structural disorder within paramyxovirus nucleoproteins and phosphoproteins. Molecular BioSystems 8, 69–81 (2012).

28. V. D. Thakkar et al., The Unstructured Paramyxovirus Nucleocapsid Protein Tail Domain Modulates Viral Pathogenesis through Regulation of Transcriptase Activity. Journal of virology, JVI.02064–02017 (2018).

29. R. M. Cox, S. A. Krumm, V. D. Thakkar, M. Sohn, R. K. Plemper, The structurally disordered paramyxovirus nucleocapsid protein tail domain is a regulator of the mRNA transcription gradient. Science Advances 3, e1602350 (2017).

30. M. Dochow, S. A. Krumm, J. E. Crowe, M. L. Moore, R. K. Plemper, Independent Structural Domains in Paramyxovirus Polymerase Protein. Journal of Biological Chemistry 287, 6878–6891 (2012).

31. S. M. Beaty et al., Efficient and Robust Paramyxoviridae Reverse Genetics Systems. mSphere 2, e00376–00316 (2017).

32. M. Habjan, N. Penski, M. Spiegel, F. Weber, T7 RNA polymerase-dependent and - independent systems for cDNA-based rescue of Rift Valley fever virus. Journal of General Virology 89, 2157–2166 (2008).

33. J. M. Emeny, M. J. Morgan, Regulation of the Interferon System: Evidence that Vero Cells have a Genetic Defect in Interferon Production. Journal of General Virology 43, 247–252 (1979).

34. U. J. Buchholz, S. Finke, K.-K. Conzelmann, Generation of Bovine Respiratory Syncytial Virus (BRSV) from cDNA: BRSV NS2 Is Not Essential for Virus Replication in Tissue Culture, and the Human RSV Leader Region Acts as a Functional BRSV Genome Promoter. Journal of Virology 73, 251–259 (1999).

35. X. Hou et al., Mutations in Sendai virus variant F1-R that correlate with plaque formation in the absence of trypsin. Medical Microbiology and Immunology 194, 129–136 (2005).

36. R. Shobana, S. K. Samal, S. Elankumaran, Prostate-specific antigen-retargeted recombinant newcastle disease virus for prostate cancer virotherapy. Journal of virology 87, 3792–3800 (2013).

