## Supplementary Figures S1-S2 and Supplementary Tables S1-S8 for "Genome-wide transposon mutagenesis of paramyxoviruses reveals constraints on genomic plasticity" (from another version of this preprint: version 1, 2020-03-31)

Title:

**Genome-wide transposon mutagenesis of paramyxoviruses reveals constraints on genomic plasticity: implications for vaccine and gene therapy**

Short title:

**Genomic scale interrogation of paramyxoviruses**

**Authors**

S. Ikegame^1†^, S. M. Beaty^1†^, C. Stevens^1^, S. T. Won^1^, A. Park^1^, D. Sachs^2^, P. Hong^1^, P. A. Thibault^1*^, B. Lee^1*^

**Affiliations**

^1^ Department of Microbiology at the Icahn School of Medicine at Mount Sinai, New York, NY 10029, USA.

^2^ Department of Genetics and Genomic Sciences at the Icahn School of Medicine at Mount Sinai, New York, NY 10029, USA.

^†^ These authors contributed equally to this work.

***Supplementary Materials***

**Supplementary Table S1.** Transposon mutagenesis calculations and metrics of SeV and MuV.

**Supplementary Figure S1.** Sequencing and transposon coverage of SeV library.

**Supplementary Table S2.** Most highly-represented insertants from SeV library.

**Supplementary Figure S2.** Sequencing and transposon coverage of MuV library.

**Supplementary Table S3.** Most highly-represented insertants from MuV library.

**Supplementary Table S4.** Transposon mutagenesis calculations and library metrics in NDV.

**Supplementary Table S5.** Most highly-represented insertants from NDV library.

**Supplementary Table S6.** Primers for generation of insertant clones.

**Supplementary Table S7.** SeV, MuV, and NDV rescue parameters.

**Supplementary Table S8.** Primers for one-step reverse-transcription and PCR of SeV, MuV, and NDV.

**Supplementary Table S1.** Transposon mutagenesis calculations and library metrics of SeV and MuV.

|  |  | | | **SeV** | **MuV** |
| --- | --- | --- | --- | --- | --- |
| **Calculations** | Genome size including eGFP gene (bp) | | | 16,278 | 16,314 |
|  | Rescue efficiency  (events per 10^5^ cells) (ref. 28) | | | 4,258 | 1,007 |
|  | Number cells per well | | | 4.0 x 10^5^ | |
|  | Number wells transfected | | | 18 | 42 |
|  | Estimated total number  of rescue events (ref. 28) | | | 3.1 x 10^5^ | 1.7 x 10^5^ |
|  | Estimated genome coverage  (rescue events per genome nt) | | | x 18.8 | x 10.4 |
| **Library Metrics** | ***Coverage:** number of nt or codons mutated/ genome size (nt) or total aa number (codons) in genome ORFs | Plasmid Library DNA (6n+18) | nt | **65.9%** (10,141 / 15,384) | **58.9%** (9,060 / 15,384) |
|  |  |  | aa | **93.4%**  (4,497 / 4,814) | **67.1%**  (3,194 / 4,760) |
|  |  | Rescued virus (P0) | nt | **50%** (7,697 / 15,384) | **20.8%** (3,191 / 15,384) |
|  |  |  | aa | **82.3%** (3,963 / 4,814) | **43.3%** (3,194 / 4,760) |
|  | Loss of coverage upon rescue (Plasmid Library minus P0) | | nt | **15.9%** | **38.1%** |
|  | **Titer** of rescued virus (iu / mL) | | | 2.2 x 10^5^ | 3.2 x 10^2^ |

* Nucleotide coverage is not >90% despite estimated 10X coverage because (a) insertions in the plasmid backbone are not accounted for, which dilutes the calculated coverage in the actual genome, (b) >1 insert per genome results in genomes with variable internal deletions that are not only non-viable, but also cannot be picked up by our sequencing library preparation and analytic pipeline, and (c) Mu-transpogenesis is known not to be completely random. For the abovementioned reasons, we carefully optimized conditions to ensure an estimated 10X coverage if feasible, while minimizing the possibility of having multiple inserts per genome. Despite the lower than expected genome-wide nucleotide coverage in the input library (6n+18), the codon-coverage is high (67.1% to 93.4%).

**Supplementary Figure S1.** Sequencing and transposon coverage of SeV library.

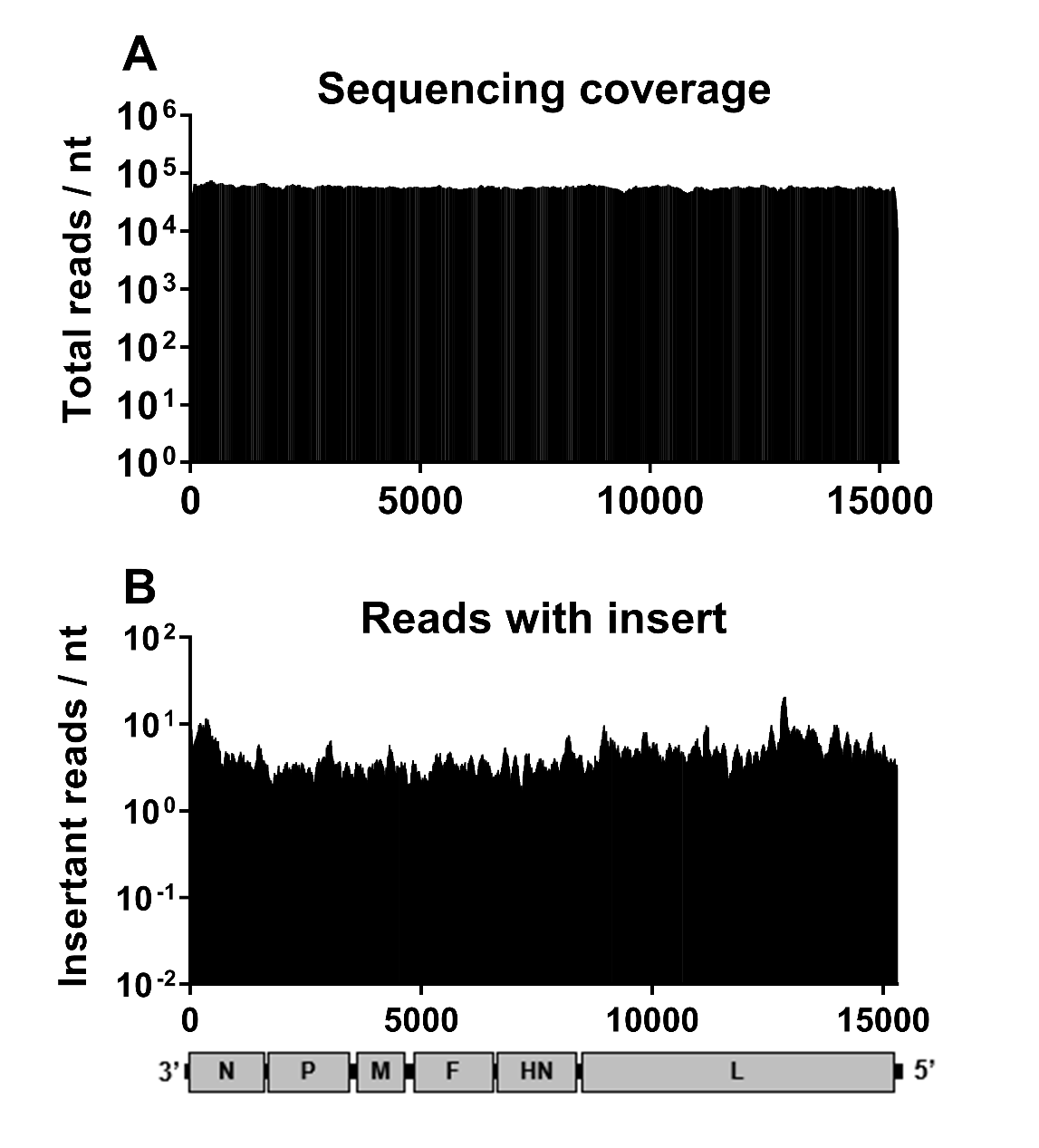

(**A**) Total number of reads at each nucleotide position in the SeV genome, regardless of transposon detection, from the input plasmid. (**B**) Distribution of insertions in a 100nt sliding window in the input plasmid DNA. To-scale schematic of the SeV genome is included at the bottom.

**Supplementary Table S2.** Most highly-represented insertants from SeV library.

|  | **Nt position in genome^a^** | **Nt count from region start** | **Avg number insertants at P2** | **Insertant nucleotide sequence^b^** | **Insertant amino acid sequence^c^** |
| --- | --- | --- | --- | --- | --- |
| **N-ORF** | **1405** | **1286** | **15** | **CTAGA\|TGCGGCCGCA\|CTAGA** | **LDAAALE** |
|  | **1606** | **1487** | **64** | **TCTGC\|TGCGGCCGCA\|TCTGC** | **SAAAASA** |
|  | 1684 | 1565 | 29 | ATAGG\|TGCGGCCGCA\|ATAGG | IGAAAIG |
| **3'UTR-N** | **1694** | **1** | **102** | **TCTAG\|TGCGGCCGCA\|TCTAG** |  |
|  | **1721** | **28** | **78** | **CTTGA\|TGCGGCCGCA\|CTTGA** |  |
|  | 1732 | 39 | 457 | TAAGA\|TGCGGCCGCA\|TAAGA |  |
| **5'UTR-P** | 1748 | 8 | 127 | GTGAA\|TGCGGCCGCA\|GTGAA |  |
|  | 1757 | 17 | 266 | CATCC\|TGCGGCCGCA\|CATCC |  |
|  | **1762** | **22** | **3848** | **ACTGA\|TGCGGCCGCA\|ACTGA** |  |
|  | 1772 | 32 | 97 | TCAGG\|TGCGGCCGCA\|TCAGG |  |
|  | 1776 | 36 | 228 | GCAAG\|TGCGGCCGCA\|GCAAG |  |
|  | 1781 | 41 | 69 | GCCAC\|TGCGGCCGCA\|GCCAC |  |
|  | 1792 | 52 | 53 | CCCCA\|TGCGGCCGCA\|CCCCA |  |
|  | 1793 | 53 | 101 | CCCAC\|TGCGGCCGCA\|CCCAC |  |
|  | **1806** | **66** | **258** | **CCCAG\|TGCGGCCGCA\|CCCAG** |  |
|  | 1807 | 67 | 61 | CCAGC\|TGCGGCCGCA\|CCAGC |  |
|  | 1813 | 73 | 87 | GTCGA\|TGCGGCCGCA\|GTCGA |  |
|  | 1814 | 74 | 130 | TCGAG\|TGCGGCCGCA\|TCGAG |  |
| **5'UTR-P**  *(C'-ORF)* | 1831 | 91 | 94 | TCGGC\|TGCGGCCGCA\|TCGGC | *SAAAASA* |
|  | 1840 | 100 | 425 | ACTTA\|TGCGGCCGCA\|ACTTA | *TLMRPQLT* |
|  | 1843 | 103 | 65 | ACCGC\|TGCGGCCGCA\|ACCGC | *TAAAATA* |
| **P-ORF**  *(C-ORF)* | **1851** | **8** | **26** | **GATCA\|TGCGGCCGCA\|GATCA** | **DHAAADQ *WIMRPQIK*** |
|  | **2007** | **164** | **22** | **ATCAA\|TGCGGCCGCA\|ATCAA** | **INAAAIN *PSMRPQST*** |
| **3'UTR-P** | 3601 | 52 | 40 | TCTAC\|TGCGGCCGCA\|TCTAC |  |
| **M-ORF** | 4696 | 1028 | 1 | ATCGG\|TGCGGCCGCA\|ATCGG | IGAAAIG |
| **F-ORF** | 6554 | 1689 | 0.7 | CTGAG\|TGCGGCCGCA\|CTGAG | AECGRTE |
| **HN-ORF** | 7168* | 476 | 1.3 | TTCTG\|TGCGGCCGCA\|TTCTG | FCAAAFW |
| **L-ORF** | 12302* | 3747 | 1.7 | CGTGG\|TGCGGCCGCA\|CGTGG | TWCGRTW |

Grey highlighted insertants were rescued and analyzed for growth. Bolded insertants were included in the competition assay.

^a^ * indicates that insertant could not be rescued.

^b^ Transposon duplicates 5nt from the site of insertion (indicated with vertical bar) and leaves a 10nt scar.

^c^ Underlined amino acids were inserted by the transposon. Italicized sequence indicates the alternative C-ORF aa sequence.

**Supplementary Figure S2.** Sequencing and transposon coverage of MuV library.

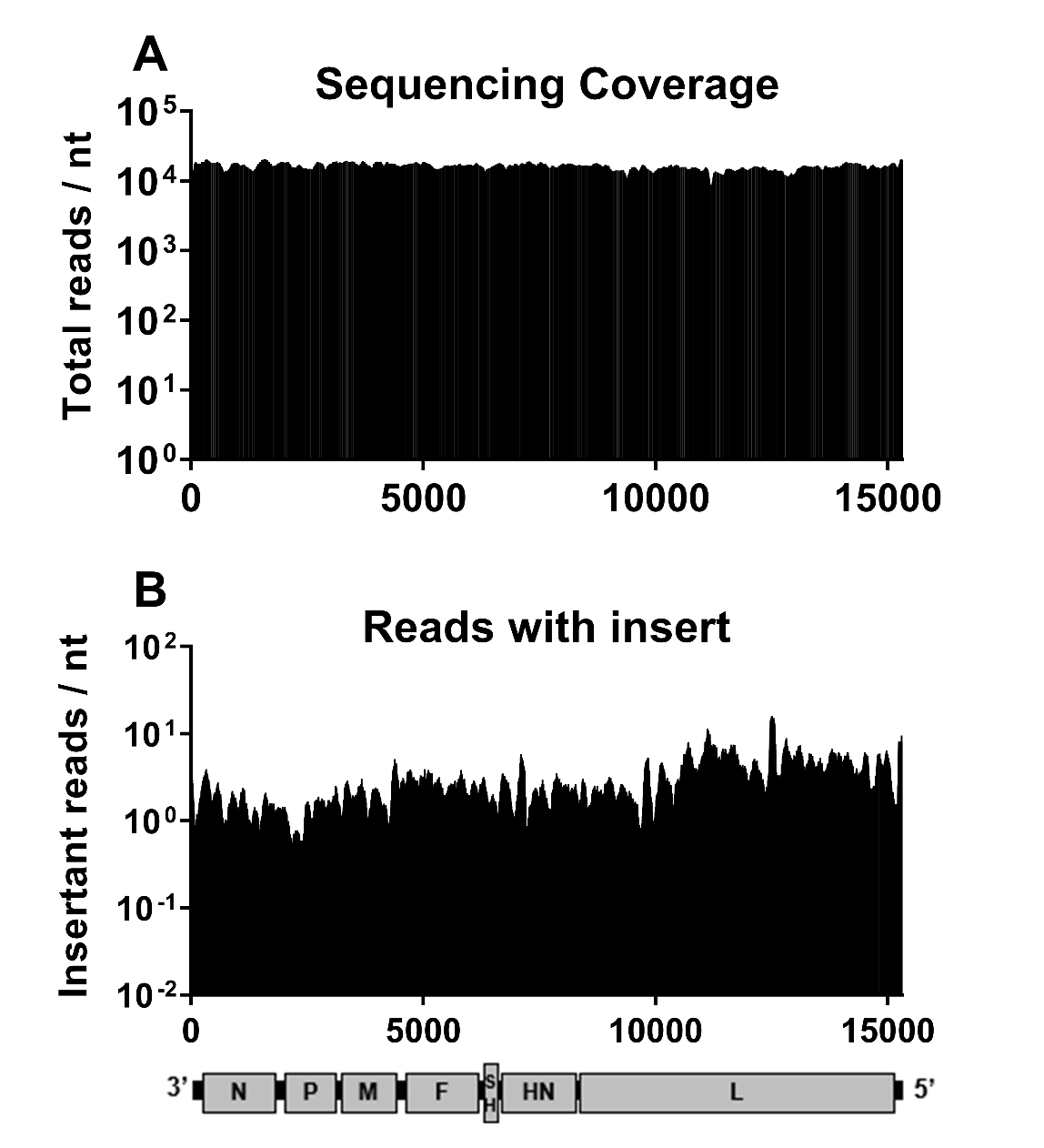

(**A**) Total number of reads at each nucleotide position in the MuV genome, regardless of transposon detection, from the input plasmid. (**B**) Distribution of insertions in a 100nt sliding window in the input plasmid DNA. To-scale schematic of the MuV genome is included at the bottom.

**Supplementary Table S3.** Most highly-represented insertants from MuV library.

|  | **Nt position in genome^a^** | **Nt count from region start** | **Avg number insertants at P2** | **Insertant nucleotide sequence^b^** | **Insertant amino acid sequence^c^** |
| --- | --- | --- | --- | --- | --- |
| **N-ORF** | 1380 | 1235 | 1 | GGTGG\|TGCGGCCGCA\|GGTGG | GGAAAGG |
|  | **1386** | **1241** | **2.7** | **CCTGT\|TGCGGCCGCA\|CCTGT** | **PVAAAPV** |
|  | **1781** | **1636** | **340** | **GGGAG\|TGCGGCCGCA\|GGGAG** | **VGVRPQGD** |
| **3'UTR-N** | 1840 | 46 | 8 | CAGGA\|TGCGGCCGCA\|CAGGA |  |
|  | **1858** | **64** | **2** | **GCTAA\|TGCGGCCGCA\|GCTAA** |  |
|  | **1881** | **87** | **21.3** | **TTCCA\|TGCGGCCGCA\|TTCCA** |  |
| **5'UTR-V/P** | 1976 | 68 | 7.3 | GCAAG\|TGCGGCCGCA\|GCAAG |  |
|  | **1977** | **69** | **1528** | **CAAGC\|TGCGGCCGCA\|CAAGC** |  |
|  | 1978 | 70 | 5.3 | AAGCC\|TGCGGCCGCA\|AAGCC |  |
| **V-ORF**  *(P-ORF)* | 1981 | 3 | 1 | CCATG\|TGCGGCCGCA\|CCATG | ()MCGRTMD |
|  | **1982** | **4** | **1** | **CATGG\|TGCGGCCGCA\|CATGG** | **()MVRPHMD** |
|  | 1984 | 6 | 1 | TGGAT\|TGCGGCCGCA\|TGGAT | MDCGRMD |
|  | **2521** | **543** | **1** | **AAGGA\|TGCGGCCGCA\|AAGGA** | **QGCGRKG** *KDAAAKE* |
| **F-ORF** | 4889* | 344 | 40 | GGTGT\|TGCGGCCGCA\|GGTGT | GVAAAGV |
|  | 4904 | 359 | 6.7 | GCACA\|TGCGGCCGCA\|GCACA | AHAAAAQ |
|  | 4905* | 360 | 1.3 | CACAA\|TGCGGCCGCA\|CACAA | AQCGRTQ |
|  | 4910 | 365 | 1 | GTGAC\|TGCGGCCGCA\|GTGAC | VTAAAVT |
| **M-ORF** | 4381 | 1118 | 2 | ATGAG\|TGCGGCCGCA\|ATGAG | MSAAAMR |
| **HN-ORF** | 7084* | 471 | 25 | TTTCT\|TGCGGCCGCA\|TTTCT | DFLRPHFS |
| **L-ORF** | 12587* | 4150 | 10 | ACCTG\|TGCGGCCGCA\|ACCTG | RPVRPQPV |

Grey highlighted insertants were rescued and analyzed for growth. Bolded insertants were included in the competition assay.

^a^ * indicates that insertant could not be rescued.

^b^ Transposon duplicates 5nt from the site of insertion (indicated with vertical bar) and leaves a 10nt scar.

^c^ Underlined amino acids were inserted by the transposon. Italicized sequence indicates the edited P-ORF aa sequence. () indicate that edited sequence contains non-coding sequence.

**Supplementary Table S4.** Transposon mutagenesis calculations and library metrics in NDV.

|  |  | | | **NDV** |
| --- | --- | --- | --- | --- |
| **Calculations** | Genome size including eGFP gene (bp) | | | 15,942 |
|  | Rescue efficiency  (events per 10^5^ cells) (ref 28) | | | 375 |
|  | Number cells per well | | | 4.0 x 10^5^ |
|  | Number wells transfected | | | 54 |
|  | Estimated total number  of rescue events (ref. 28) | | | 8.1 x 10^4^ |
|  | Estimated genome coverage  (rescue events per genome nt) | | | x 5.1 |
| **Library Metrics** | **Coverage:** number of nt or codons mutated/ genome size (nt) or total aa number (codons) in genome ORFs | Plasmid DNA (6n+18) | nt | **51.5%** (7,792 / 15,130) |
|  |  |  | aa | **78.6%**  (3,606 / 4,588) |
|  |  | Rescued virus (P0) | nt | **34.4%** (5,205 / 15,130) |
|  |  |  | aa | **61.5%** (2,820 / 4,588) |
|  | Loss of coverage upon rescue (Plasmid Library minus P0) | | nt | **17.1%** |
|  | **Titer of rescued virus (iu / mL)** | | | 1.0 x 10^1^ |

**Supplementary Table S5.** Most highly-represented insertants from NDV library.

|  | **Nt position in genome^a^** | **Nt count from region start** | **Average number of insertants at P2** | **Insertant nucleotide sequence^b^** | **Insertant amino acid sequence^c^** |
| --- | --- | --- | --- | --- | --- |
| **F-ORF** | **5367** | **880** | **49** | **CGGGA\|TGCGGCCGCA\|CGGGA** | **GMRPHGNL** |
|  | 5383 | 896 | 442 | ATGCG\|TGCGGCCGCA\|ATGCG | MRAAMRA |
|  | **5384** | **897** | **40** | **TGCGT\|TGCGGCCGCA\|TGCGT** | **RCGRMRA** |
| **L-ORF** | 11643 | 3319 | 93 | CTTAG\|TGCGGCCGCA\|CTTAG | LVRPHLVS |
|  | 11867 | 3543 | 48 | ATGAC\|TGCGGCCGCA\|ATGAC | DCGRNDT |
|  | 11868 | 3544 | 122 | TGACA\|TGCGGCCGCA\|TGACA | DMRPHDTS |
|  | 11869 | 3545 | 281 | GACAC\|TGCGGCCGCA\|GACAC | DTAAADTS |
|  | 11870* | 3546 | 8857 | ACACC\|TGCGGCCGCA\|ACACC | TCGRNTS |
|  | 11871 | 3547 | 830 | CACCA\|TGCGGCCGCA\|CACCA | TMRPMTSK |
|  | 11872* | 3548 | 65000 | ACCAG\|TGCGGCCGCA\|ACCAG | TSAAATSK |
|  | ***11872^#^***  ***+ F^rev^*** | ***3548*** | ***65000*** | ***ACCAG\|TGCGGCCGCA\|ACCAG*** | ***TSAAATSK*** |
|  | 11873 | 3549 | 868 | CCAGC\|TGCGGCCGCA\|CCAGC | SCGRTSK |
|  | 11874* | 3550 | 810 | CAGCA\|TGCGGCCGCA\|CAGCA | SMRPHSKN |
|  | 11875 | 3551 | 727 | AGCAA\|TGCGGCCGCA\|AGCAA | SNAAASKN |
|  | 11876 | 3552 | 269 | GCAAG\|TGCGGCCGCA\|GCAAG | KCGRSKN |
|  | 11877 | 3553 | 84 | CAAGA\|TGCGGCCGCA\|CAAGA | KMRPHKN |
|  | 12416 | 4092 | 49 | TTGGG\|TGCGGCCGCA\|TTGGG | GCGRIGV |
|  | 12727 | 4403 | 44 | GGAGC\|TGCGGCCGCA\|GGAGC | AAAAAAAL |
|  | 13213 | 4889 | 192 | CCATT\|TGCGGCCGCA\|CCATT | PFAAAPLL |
|  | 13215* | 4891 | 3286 | ATTAC\|TGCGGCCGCA\|ATTAC | LLRPQLL |
|  | 14818 | 6494 | 34 | ATTAC\|TGCGGCCGCA\|ATTAC | ITAAAIT |

Grey highlighted insertants were rescued and analyzed for growth.

Bold insertants demonstrated syncytia and continuous GFP spread after rescue.

^a^ * indicates that insertant did not demonstrate syncytia or GFP spread after rescue.

^#^ indicates the insertant rescued with the fusion-revertant compensatory point mutations

^b^ Transposon duplicates 5nt from the site of insertion (indicated with vertical bar) and leaves a 10nt scar.

^c^ Underlined amino acids were inserted by the transposon.

**Supplementary Table S6.** Primers for generation of insertant clones.

| **Virus** | **Insertant** | **Primer sequence (5' to 3')** |
| --- | --- | --- |
| **SeV** | 1405-N | For: TGCGGCCGCACTAGAAACAGAAGCTCATGCGGAC |
|  |  | Rev: TCTAGTGCGGCCGCATCTAGGTCGATATCGGCATTG |
|  | 1606-N | For: TGCGGCCGCATCTGCAACCCATGGAGATGAAGGCCGC |
|  |  | Rev: GCAGATGCGGCCGCAGCAGAATCCTCTTGCCGT |
|  | 1684-N | For: TGCGGCCGCAATAGGAGGAATCTAGGATCATACGAGG |
|  |  | Rev: CCTATTGCGGCCGCACCTATCCCAGCTACTGCTG |
|  | 1694-UTR-N | For: TGCGGCCGCATCTAGGATCATACGAGGCTTCAAGGTAC |
|  |  | Rev: CTAGATGCGGCCGCACTAGATTCCTCCTATCCCAGCT |
|  | 1721-UTR-N | For: TGCGGCCGCACTTGATCCGTAGTAAGAAAAACTTAGGGT |
|  |  | Rev: TCAAGTGCGGCCGCATCAAGTACCTTGAAGCCTCGTA |
|  | 1762-UTR-N | For: TGCGGCCGCAACTGATCGGCTCAGGCAAGGCCACA |
|  |  | Rev: TCAGTTGCGGCCGCATCAGTGGATGAACTTTCACCC |
|  | 1772-UTR-P | For: TGCGGCCGCATCAGGCAAGGCCACACCCAAC |
|  |  | Rev: CCTGATGCGGCCGCACCTGAGCCGATCAGTG |
|  | 1806-UTR-P | For: TGCGGCCGCACCCAGCAGTCGAGACAGCCAC |
|  |  | Rev: CTGGGTGCGGCCGCACTGGGTGTGGTCGGTG |
|  | 1840-UTR-P | For: TGCGGCCGCACTTACCGCATGGATCAAGATGCCTT |
|  |  | Rev: GTAAGTGCGGCCGCAGTAAGTGTAGCCGAAGCC |
|  | 1851-P | For: TGCGGCCGCAGATCAAGATGCCTTCATTCTTAAAGAAGATTCT |
|  |  | Rev: TGATCTGCGGCCGCATGATCCATGCGGTAAGTGTAG |
|  | 2007-P | For: TGCGGCCGCAATCAACACTCCCCAAGGACCA |
|  |  | Rev: TTGATTGCGGCCGCATTGATGGTGTTGTGGAGCC |
|  | 4696-M | For: TGCGGCCGCAATCGGAAGGATCAGAAAGCTGTAAAT |
|  |  | Rev: CCGATTGCGGCCGCACCGATGTTCTTAGCCACAACA |
|  | 6554-F | For: TGCGGCCGCACTGAGAAAAGATGATCACGACCATTAT |
|  |  | Rev: CTCAGTGCGGCCGCACTCAGCCATCGCATCAAACC |
|  | 7168-HN | For: TGCGGCCGCATTCTGGAGATGCCCTGTCGGAGAAC |
|  |  | Rev: CAGAATGCGGCCGCACAGAAACTATGTGGCTCAAGTG |
|  | 12302-L | For: TGCGGCCGCACGTGGGCCTACGGGACTGATGAGAT |
|  |  | Rev: CCACGTGCGGCCGCACCACGTATACACCATAGCTATC |
| **MuV** | 1386-N | For: TGCGGCCGCACCTGTGAATCCTTTTGTTCCTCCGGTT |
|  |  | Rev: ACAGGTGCGGCCGCAACAGGCCCACCTGGTATCC |
|  | 1781-N | For: TGCGGCCGCAGGGAGACTGGGATGAGTAAATCACTGACA |
|  |  | Rev: CTCCCTGCGGCCGCACTCCCACTTGCAACTGTGCGTT |
|  | 1858-UTR-N | For: TGCGGCCGCAGCTAACTGCCCAAATCCACTACATTCCA |
|  |  | Rev: TTAGCTGCGGCCGCATTAGCTGTGGCTGGATTGTCCT |
|  | 1881-UTR-N | For: TGCGGCCGCATTCCATTCATATTTAGTCTTTAAGAAA |
|  |  | Rev: TGGAATGCGGCCGCATGGAATGTAGTGGATTTGGGC |
|  | 1976-UTR-P | For: TGCGGCCGCAGCAAGCCATGGATCAATTTATAAAACA |
|  |  | Rev: CTTGCTGCGGCCGCACTTGCCCGGAAAGAAACACG |
|  | 1977-UTR-P | For: TGCGGCCGCACAAGCCATGGATCAATTTATAAAACAGGA |
|  |  | Rev: GCTTGTGCGGCCGCAGCTTGCCCGGAAAGAAACACG |
|  | 1982-P | For: TGCGGCCGCACATGGATCAATTTATAAAACAGGATGAG |
|  |  | Rev: CCATGTGCGGCCGCACCATGGCTTGCCCGGAA |
|  | 2521-P | For: TGCGGCCGCAAAGGAGAGGTCCGGGTCTTTGAGTG |
|  |  | Rev: TCCTTTGCGGCCGCATCCTTGGACCCAGCTGAGG |
|  | 4381-M | For: TGCGGCCGCAATGAGAAACCTATGAGAATTTCCTCTA |
|  |  | Rev: CTCATTGCGGCCGCACTCATTGAGGCAGACTTCCTG |
|  | 4889-F | For: TGCGGCCGCAGGTGTTGCGACCGCAGCACAAGTGA |
|  |  | Rev: ACACCTGCGGCCGCAACACCGAGCGCAGCAATG |
|  | 4905-F | For: TGCGGCCGCACACAAGTGACTGCCGCTGTCTCATT |
|  |  | Rev: TTGTGTGCGGCCGCATTGTGCTGCGGTCGCAA |
|  | 7084-HN | For: TGCGGCCGCAATTTCTCCATCGGCCATCCACTTAACAT |
|  |  | Rev: GAAATTGCGGCCGCAGAAATCATGGGTTGCATAATCTTC |
|  | 12587-L | For: TGCGGCCGCAACCTGTCGAGTCTTGTGTAGTAAATCCGC |
|  |  | Rev: CAGGTTGCGGCCGCACAGGTCTTATACAGCAAGATGAGC |
| **NDV** | 5367-F | For: TGCGGCCGCACGGGAACCTAAATAATATGCGTGCCACCTAC |
|  |  | Rev: TCCCGTGCGGCCGCATCCCGACTGAAGGTAGAGTTACC |
|  | 5383-F | For: TGCGGCCGCAATGCGTGCCACCTACTTGGAAACCTTATCC |
|  |  | Rev: CGCATTGCGGCCGCACGCATATTATTTAGGTTCCCGACTGA |
|  | 11870-L | For: TGCGGCCGCAACACCAGCAAGAATCCTCCGATGAGGGT |
|  |  | Rev: GGTGTTGCGGCCGCAGGTGTCATCGGTCAATTCTATATTGC |
|  | 11872-L | For: TGCGGCCGCAACCAGCAAGAATCCTCCGATGAGGGTAC |
|  |  | Rev: CTGGTTGCGGCCGCACTGGTGTCATCGGTCAATTCTATA |
|  | 11874-L | For: TGCGGCCGCACAGCAAGAATCCTCCGATGAGGGTACCA |
|  |  | Rev: TGCTGTGCGGCCGCATGCTGGTGTCATCGGTCAATTCTA |
|  | 13215-L | For: TGCGGCCGCAATTACTTAGCCCCGATCAAGTGAGCTC |
|  |  | Rev: GTAATTGCGGCCGCAGTAATGGTTTCACAGCATCCGACAG |

**Supplementary Table S7.** SeV, MuV, and NDV rescue transfection parameters.

|  | **SeV** | **MuV** | **NDV** |
| --- | --- | --- | --- |
| Genome (μg) | 4.00 | 5.00 | 0.8 |
| pTM1-N (μg) | 1.44 | 0.30 | 0.4 |
| pTM1-P (μg) | 0.77 | 0.10 | 0.2 |
| pTM1-L (μg) | 0.07 | 0.20 | 0.2 |
| pCAGGS-T7opt (μg) | 4.00 | 2.00 | 0.4 |
| OptiMEM (μL) | 200.00 | 200.00 | 200.0 |
| Plus reagent (μL) | 5.50 | 7.50 | 2.0 |
| Lipofectamine LTX (μL) | 8.90 | 18.75 | 5.0 |

**Supplementary Table S8.** Primers for one-step reverse-transcription and PCR of SeV, MuV, and NDV.

| **Pair** | | **Forward** | **Reverse** |
| --- | --- | --- | --- |
| **SeV** | 1 | ACCAAACAAGAGAAAAAACATGTATGGAAT | GGTCACGAACTCCAGCAG |
|  | 2 | AAGCGCGATCACATGGTC | TCCCGACCTTTCTCCTGAT |
|  | 3 | TGGTGCACCTCGGGTTG | CATGGTGGACTGCGATCG |
|  | 4 | CACTCAGCTCTGTGAGAGTA | GTATAGTTGTCTACAGCACATTCATAAG |
|  | 5 | AGGGTCCAATACGGCAATCT | GTTATTAAGATGTAAGTGCAATATCAGTG |
|  | 6 | GAAAGGTTCCTTAGGGAAGC | ACCAGACAAGAGTTTAAGAGATATTTATTC |
| **MuV** | 1 | ACCAAGGGGAGAATGAATATGGGATATTG | ATGTGATCGCGCTTCTCGTTGG |
|  | 2 | TCCGCCCTGAGCAAAGACC | CGAATTACCACCGGTCAAATTTGCTTGA |
|  | 3 | CCACTGATGCCTATAGGAGAATCAACAA | GAAAAGGATGGAATTCTCGTGCAACC |
|  | 4 | CCCACTGCAACCTCACCCAAT | CCAGTCTTGCCTTGGACAGCTTATT |
|  | 5 | GATCTCAGCATATTCATGAAGGATAAGGC | GATGAGCCTGTGTGCAAGTGTAATG |
|  | 6 | CTCCAATCAACTTCTCCGTTTATGAAACCA | ACCAAGGGGAGAAAGTAAAATCAATTTTTTCTTAAAAGG |
| **NDV** | 1 | ACCAAACAGAGAATCCGTGAGTTACG | GTTGGATTTCGGACCGCATCATAG |
|  | 2 | GTCTTGAAACAGACATCCTCCATCC | GTGTCATGATGATCTGGGTGAGTG |
|  | 3 | TCTGAGATTGCGCTCCGCC | TCCGGAGCACACCAAGTGCTAAAT |
|  | 4 | GATGCAGAGATCACTCACATTCATATCAGT | CGCCTAAACGAGGCAAGCCAATTA |
|  | 5 | GCAATCGCACACCCCAACGA | ACCGCAACAGGTGCTTCTCTGAT |
|  | 6 | CACTGCACCTACATAGTAAATTTAGTTGCTGT | ACCAAACAAAGATTTGGTGAATGACGAGAC |
